# Intra-cluster receptor density (IRD): a molecular switch for TNFR1 clusters’ signaling

**DOI:** 10.1101/2024.08.09.607302

**Authors:** Subhamoy Jana, Priyanka Roy, Jibitesh Das, Basab Bijayee Bera, Parijat Biswas, Nandana Nanda, Bidisha Sinha, Deepak Kumar Sinha

**Author notes:** Corresponding author: Deepak Kumar Sinha. Equal first-author contribution (Subhamoy Jana, Priyanka Roy).

## Abstract

Tumor Necrosis Factor Receptor 1 (TNFR1) signaling regulates cell fate in inflammation, immune responses, and tumorigenesis. While TNF-α–mediated TNFR1 pathways are well known, the role of receptor clustering remains unclear. Utilizing homo-FRET using fluorescence anisotropy, we show that intra-cluster receptor density (IRD) governs TNFR1 signaling outcomes. Soluble TNF-α (sTNF-α) increases IRD at cluster cores but decreases it at rims via receptor reorganization. Reducing IRD through membrane tension, zafirlukast, actin depolymerization, or cholesterol depletion suppresses sTNF-α signaling, whereas increasing IRD by lowering membrane tension or exposing cells in a 3D gel-like microenvironment triggers ligand-independent activation. These findings reveal IRD as a key regulator of receptor signaling, with potential relevance across related receptor families and innovative strategies in modulating TNFR1 signaling.

**Teaser:** Intra-cluster receptor density (IRD) is critical for TNFR1 signal modulation, with higher IRD activating and lower IRD impairing the TNFR1 signaling.

## Introduction

Tumor necrosis factor receptor 1 (TNFR1) is a type I transmembrane protein of the TNF receptor superfamily (TNFRSF) (*1, 2*), which plays a multifaceted role in modulating numerous cellular processes, encompassing inflammation (*3, 4*), cell survival (*5*), apoptosis (*6, 7*), and differentiation (*8*). While TNF-α-mediated TNFR1 signaling pathways (*9, 10*) are widely known, the role of receptor clustering in downstream signal activation remains unexplored. Studies in cell-free systems suggest that the trimeric ligand TNF-α (*11*) interacts with TNFR1s in their native monomeric and/or dimeric state (*12*) to form the trimeric TNFR1 primary cluster(*13*) , which acts as the basic signaling unit. Further, the interaction among pre-ligand assembly domains (PLAD) across different TNFR1 primary clusters promotes the formation of ligand-independent, higher-order TNFR1 secondary clusters on the cell membrane (*14–19*). Besides PLAD, TNFR1 interacts via its other domains, including cysteine-rich domains (CRDs) (*20*), transmembrane domain (TMD) (*21*), and cytoplasmic domain (CD) (*22, 23*), assembling ligand-receptor and receptor-receptor complexes into larger clusters. Our observation, along with others, indicates that TNFR1 consistently forms pre-formed clusters on the cell membrane even without additional ligand stimulation (*18, 22*).

Understanding the importance of TNFR1 clustering in downstream signaling is crucial for therapeutic advancements. The targeted treatment using small molecule inhibitors (**table S1**) and/or the polymorphism-induced TNFR1 dysfunctions (*24–29*) is limited, primarily focusing on ligand sequestration or decoy receptors (*30*). Thus, it highlights the need for a deeper understanding of TNFR1 cluster organization and its role in modulating downstream signaling across various conditions.

TNFR1 clustering has been studied with its ligand at basal and stimulated states (*14*), but the organization within pre-formed clusters and their impact on downstream signaling remain unclear. A previous study on TNFR1 signal activation by reinforcing secondary clusters with agonistic antibodies (*31, 32*) suggests that an increase in receptor density within the cluster activates the signaling. In our study, we showed that the density of TNFR1 receptors within pre-formed TNFR1 clusters, referred to as intra-cluster receptor density (IRD), is a crucial determinant of its downstream signaling. We anticipated that IRD modulates signaling efficacy through cooperativity. In this context, cooperativity refers to a collective behavior in which receptor activation is influenced by the spatial proximity of neighboring receptors within a cluster, thereby modulating its likelihood of activation.

Manipulating the IRD while preserving other biochemical parameters, such as chemical composition and the biomolecular organization, is a challenging task. Using HeLa as a model system, we subjected cells to diverse conditions that alter the TNFR1 cluster’s IRD to examine its link with the TNFR1 downstream signaling state. Initially, we investigated whether the activation of TNFR1 signaling by soluble TNF-α (sTNF-α) or inhibition by the antagonist zafirlukast (*33, 34*) occur by modulating the IRD. Further, we explored other avenues that could influence IRD, namely, altered plasma membrane tension, actin depolymerization, changes in extracellular microenvironment, and cholesterol depletion on TNFR1’s IRD and signaling efficacy. We used the fluorescence anisotropy-based homo-FRET technique to assess TNFR1 IRD, where loss of anisotropy reflects increased homo-FRET (*35–38*) and receptor proximity and thus, IRD. Variations in fluorescence anisotropy within TNFR1-EGFP clusters indicate differential homo-FRET efficiency, where the anisotropy map inversely correlates with the TNFR1 IRD, as elucidated in Materials and Methods.

The canonical NF-κB pathway is a major signaling cascade activated downstream of TNFR1, promoting pro-inflammatory and pro-survival responses. Upon ligand binding, TNFR1 recruits adaptor proteins, such as TNF Receptor Type 1-Associated Death Domain Protein (TRADD), TNF Receptor–Associated Factor 2 (TRAF2), Receptor-Interacting Protein Kinase 1(RIPK1), and Cellular Inhibitor of Apoptosis Protein 1 and 2 (cIAP1/2), via its cytoplasmic death domain to form the membrane-bound Complex I. A critical step in this pathway is the ubiquitination of RIPK1 by cIAP1/2, which attaches K63- and M1-linked ubiquitin chains. These chains act as platforms for the recruitment of downstream signaling complexes, including Linear Ubiquitin Chain Assembly Complex (LUBAC), TAK1-Binding Protein 2 and 3 (TAB2/3), Transforming Growth Factor Beta– Activated Kinase 1 (TAK1), and the IκB Kinase Complex (IKK) complex (composed of IKKα, IKKβ, and NF-κB Essential Modulator (NEMO)) (*39–41*). TNFR1 signal activation triggers IKK phosphorylation, leading to Nuclear Factor Kappa-Light-Chain-Enhancer of Activated B Cells (NF-κB (p65/p50)) nuclear translocation (*41*). We assessed the downstream signaling of TNFR1 by NF-κB activation assay in different physiological conditions, which can alter the TNFR1 IRD.

This manuscript demonstrates that altering the TNFR1 IRD changes the response to TNF-α in downstream signaling, both in a ligand-dependent and ligand-independent manner. Our findings provide valuable insights into the role of clustering in TNFRSF function and its potential implications for therapeutic strategies.

## Results

### Fluorescence anisotropy mapping reveals heterogeneous packing in TNFR1 clusters

We cloned human TNFR1 into the mEGFP-N1 backbone to generate a TNFR1-EGFP construct (**Fig. 1A**). Immunostaining in transiently transfected HeLa cells confirmed that TNFR1-EGFP assembled into membrane clusters resembling endogenous TNFR1. We designated clusters lacking EGFP as “endogenous” (**fig. S1A**) and those containing EGFP as “exogenous” (**Fig. 1B and fig. S1B**). Time-lapse imaging at 20 Hz in Total Internal Reflection Fluorescence (TIRF) mode revealed exponential photobleaching of exogenous clusters, consistent with a large number of TNFR1-EGFP molecules per cluster (**fig. S1C**). In contrast, clusters composed of only a few molecules are expected to photo-bleach in discrete steps (*42–44*). The colocalization of EGFP and anti-TNFR1 (**Fig. 1B and fig. S1B**), together with sequencing validation, confirmed the construct’s accuracy. The fraction of TNFR1-EGFP clusters localized to the plasma membrane increased with time after transfection (**fig. S1D**) reaching ∼76 ± 8% by 18 hours (**Fig. 1C and movie S1**). A minority of clusters displayed high mobility (**movie S2**), likely reflecting intracellular vesicles. Treatment with the dynamin inhibitor dynasore (*45*) reduced the fraction of mobile clusters (**movie S3**). However, a considerable number of mobile clusters remained, suggesting that they may arise from multiple sources, including endocytic vesicles as well as transport vesicles carrying TNFR1-EGFP from the ER, through the Golgi, to the plasma membrane.

**Fig. 1.**
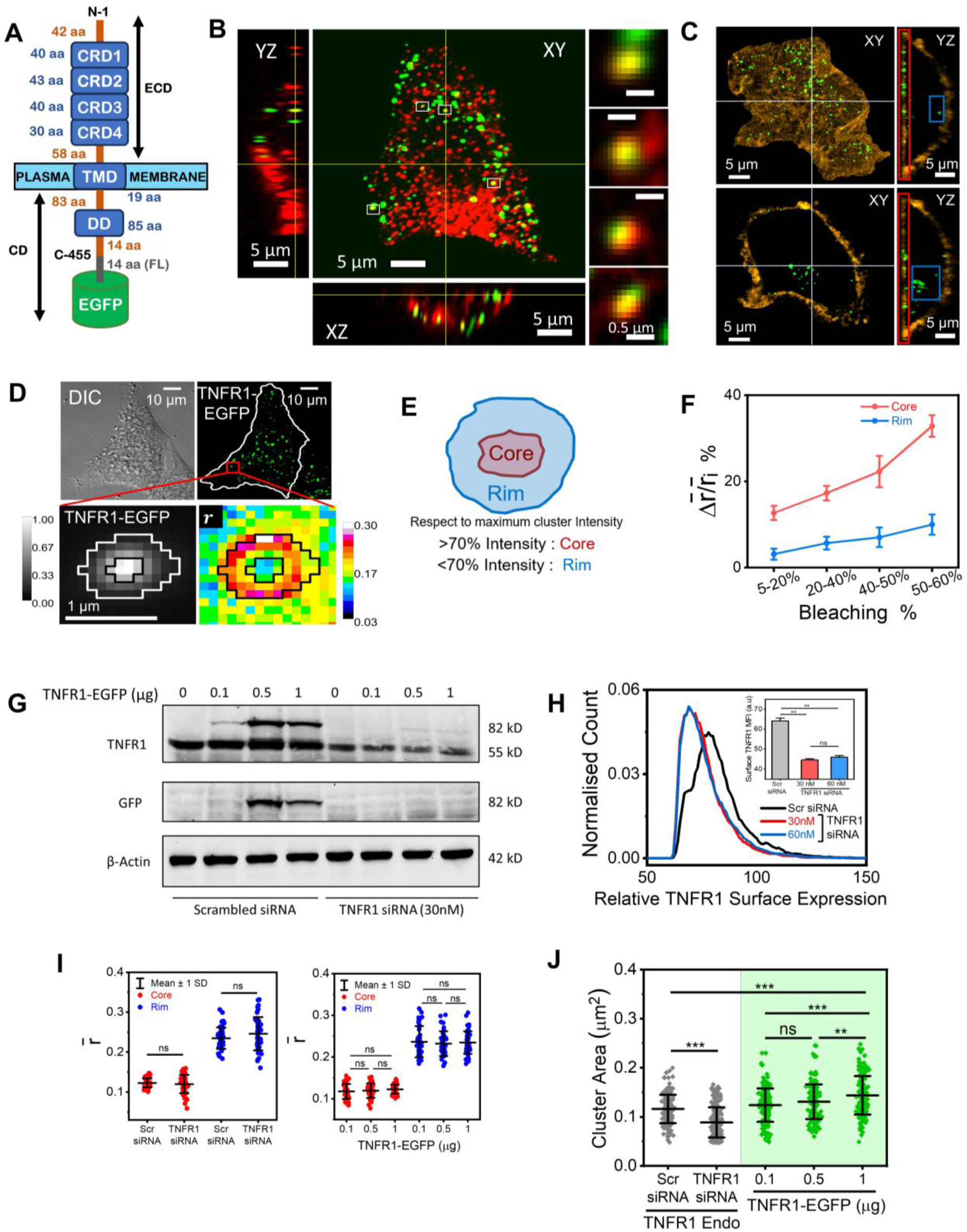
Homo-FRET–based mapping of intra-cluster receptor density (IRD) in TNFR1-EGFP nanoclusters. (**A**) Schematic representation of the TNFR1-EGFP fusion construct showing the extracellular domain (ECD), cysteine-rich domains (CRD1–4), transmembrane domain (TMD), cytoplasmic domain (CD), and death domain (DD). An additional 14-amino-acid (aa) flexible linker (FL) connects the C terminus of TNFR1 to EGFP. The amino acid lengths of individual domains are indicated. (**B**) Confocal images of a HeLa cell expressing TNFR1-EGFP (green) and immunostained with anti-TNFR1 antibody (red) in XY, YZ, and XZ planes. Merged and magnified views of four representative clusters demonstrate colocalization between TNFR1-EGFP and endogenous TNFR1. Images were processed to reduce noise by Gaussian averaging (radius = 1) using ImageJ. (**C**) Representative confocal Airyscan images showing orthogonal sections (XY and YZ) of a HeLa cell expressing TNFR1-EGFP (green) and labeled with CellMask Orange (orange) to visualize the plasma membrane. Basal (z = 0; top) and mid-plane (z = 10; bottom) sections highlight TNFR1 clusters localized at the plasma membrane (red box) and within the cytoplasm (blue box). (**D**) Differential interference contrast (DIC; top left) and epifluorescence (top right) images of a TNFR1-EGFP–expressing HeLa cell. Enlarged views display intensity (bottom left) and fluorescence anisotropy (bottom right) maps of a representative TNFR1-EGFP cluster. (**E**) Schematic illustration of a TNFR1 cluster depicting two sub-regions: a densely packed central core and a peripheral region of lower receptor density. (**F**) Percentage change in relative mean fluorescence anisotropy (Δ 𝑟̅⁄𝑟̅_𝑖_) plotted against the percentage of photobleaching for the core (red) and rim (blue) regions of TNFR1–EGFP clusters (n = 20) in live HeLa cells. Here, Δ𝑟̅ = 𝑟_𝑓̅_ − 𝑟̅_𝑖_where 𝑟_𝑓̅_ and 𝑟̅_𝑖_represent post- and pre-bleach mean anisotropy, respectively. Data are means ± SEM from 3 cells each across two independent experiments. (**G**) Immunoblot analysis of HeLa cells transfected with scrambled or TNFR1 siRNA and co-transfected with increasing doses (0-1 μg) of TNFR1-EGFP. Blots were probed with anti-TNFR1 and anti-GFP antibodies; β-Actin served as a loading control. Representative of two independent experiments. (**H**) Flow cytometric analysis of surface TNFR1 levels in control (scrambled siRNA) and TNFR1 siRNA–treated HeLa cells (30 nM and 60 nM, day 4). The inset shows quantification of median fluorescence intensity (MFI). Representative of two independent experiments. (**I**) Mean fluorescence anisotropy (𝑟̅) of core (red) and rim (blue) regions of TNFR1-EGFP clusters (n = 45) in HeLa cells transfected with scrambled siRNA (Scr siRNA) or TNFR1 siRNA (left), and in cells overexpressing with increasing doses (0-1 μg) of TNFR1-EGFP (right). The data represent 3 cells each from three independent experiments. **(J)** Quantification of cluster area (μm^2^) for endogenous TNFR1 clusters (n ≥ 170) in scrambled siRNA (Scr siRNA) and TNFR1 siRNA-treated HeLa cells (left, gray), and for exogenous TNFR1-EGFP clusters (n = 170) in cells expressing different doses (0-1 μg) of TNFR1-EGFP (right, green), acquired by confocal Airyscan imaging. The data represent 8 cells each from three independent experiments. Statistical significance between datasets was determined by one-way ANOVA (H-J) and is indicated in the figures as follows: ∗P < 0.05; ∗∗P < 0.01; ∗∗∗P < 0.001; ns, not significant. Scale bars, as indicated in the corresponding images.

Fluorescence anisotropy (𝑟) mapping of TNFR1-EGFP clusters in live cells revealed marked intra-cluster heterogeneity (**Fig. 1D**). The central core exhibited lower anisotropy (𝑟_𝑐̅𝑜𝑟𝑒_) than the outer rim (𝑟̅_𝑟𝑖𝑚_) (**Fig. 1E**), suggesting that the intra-cluster variations in 𝑟 could originate from differing homo-FRET efficiencies. The lower 𝑟_𝑐̅𝑜𝑟𝑒_ value implies a heightened homo-FRET efficiency in the core, indicating a denser packing of TNFR1 molecules in the core than in the rim or a higher IRD in the core than in the rim. The anisotropy reflects the degree of polarization of the fluorescence, and it decreases when emitted photons get depolarized due to factors such as molecular rotation. Homo-FRET, a fluorescence resonance energy transfer between identical fluorophores in close proximity, leads to such depolarization without requiring molecular rotation (*36*). This occurs because the energy is transferred to a neighboring fluorophore with an unfavorably oriented emission dipole, which can be established using EGFP dimers. Thus, homo-FRET is expected to cause a measurable loss in anisotropy, making anisotropy a sensitive readout for detecting molecular proximity and clustering of identical fluorophores. We validated this relationship by comparing monomeric EGFP with EGFP dimers (2EGFP), where photobleaching-induced conversion of dimers to monomers increased anisotropy (**fig. S1E**) (*46*). A similar anisotropy increase occurred in TNFR1-EGFP clusters upon photobleaching (**fig. S1F**), confirming the homo-FRET as the origin of intra-cluster anisotropy variations. Photobleaching produced a greater anisotropy increase in the core than in the rim (**Fig. 1F**), consistent with denser packing in the core (*37, 38*). When homo-FRET was nearly eliminated, anisotropy maps became uniform (**fig. S1G**), excluding the possibility of imaging or computational artifacts. To test whether higher EGFP mobility at the rim influenced anisotropy, we varied linker length, but observed minimal changes in 𝑟_𝑐̅𝑜𝑟𝑒_or 𝑟̅_𝑟𝑖𝑚_(**fig. S2A**), reinforcing that anisotropy differences arise from IRD variation rather than tag flexibility. Therefore, we used a 14 aa linker-length plasmid of TNFR1-EGFP in all subsequent experiments unless specified otherwise.

We next altered receptor abundance by depleting cellular TNFR1 levels. Optimal TNFR1 knockdown was achieved with 30 nM siRNA over 4 days (**fig. S2B**). Immunoblotting confirmed dose-dependent TNFR1-EGFP expression, plateauing at 0.5 μg plasmid **(Fig. 1G**). Knockdown reduced both endogenous and exogenous TNFR1 (**Fig. 1G**), including ∼30% of endogenous surface receptors **(Fig. 1H**). Fluorescence anisotropy measurement revealed that TNFR1 depletion or overexpression did not alter TNFR1-EGFP clusters’ IRD (**Fig. 1I**). However, depletion significantly reduced endogenous cluster area (**Fig. 1J**), and overexpression of TNFR1-EGFP did not rescue the endogenous cluster size (**fig. S2C**). Instead, overexpression increased the size of exogenous clusters in a dose-dependent manner (**Fig. 1J**), while endogenous clusters remained surprisingly unaffected (**fig. S2C**). In siRNA-treated cells, TNFR1-EGFP cluster areas failed to scale with plasmid dose and remained smaller than in controls (**fig. S2D**). Importantly, 𝑟_𝑐̅𝑜𝑟𝑒_ and 𝑟̅_𝑟𝑖𝑚_ values were independent of cluster area across all conditions (**fig. S2E**).

### Dynamic and spontaneous rearrangement of receptor molecules occurs within TNFR1 clusters

We examined the intra-cluster dynamics of TNFR1 molecules under physiological conditions. Time-lapse anisotropy maps of TNFR1-EGFP clusters in live HeLa cells (**Fig. 2A**) revealed significant temporal variations in 𝑟_𝑐̅𝑜𝑟𝑒_. To ascertain that the temporal variations in 𝑟_𝑐̅𝑜𝑟𝑒_ result from dynamic changes in local IRD within the core and not instrumental noise, we compared the changes in the cluster’s 𝑟_𝑐̅𝑜𝑟𝑒_in live cells with those in fixed cells (**Fig. 2B**), and observed a significantly smaller temporal variation (**Fig. 2, C and D**). We hypothesized that the crosslinking of proteins hinders the intra-cluster reorganization of TNFR1-EGFP molecules. The notably lower variations (±5%) of 𝑟_𝑐̅𝑜𝑟𝑒_in fixed clusters compared to live clusters (±20%) indicated that the temporal variations of 𝑟_𝑐̅𝑜𝑟𝑒_in live cells, originated from a change in the local IRD due to intra-cluster reorganization of receptors. The anisotropy changes in the fixed cluster (**Fig. 2D)** further suggest that a change below 5% in our experimental setup could arise from measurement noise. A lack of fluorescence recovery after photobleaching (FRAP) of the TNFR1-EGFP cluster (**Fig. 2E**) established a negligible change in the total number of receptor molecules associated with the cluster within 90-second. Furthermore, the temporal variations of 𝑟_𝑐̅𝑜𝑟𝑒_ and 𝑟̅_𝑟𝑖𝑚_were consistently higher in live cells compared to fixed cells, regardless of the TNFR1-EGFP linker flexibility (**fig. S3**). This suggests that the increased temporal fluctuations in anisotropy originate from dynamic intra-cluster reorganization of the receptor, rather than from enhanced EGFP mobility due to linker flexibility in live cells.

**Fig. 2.**
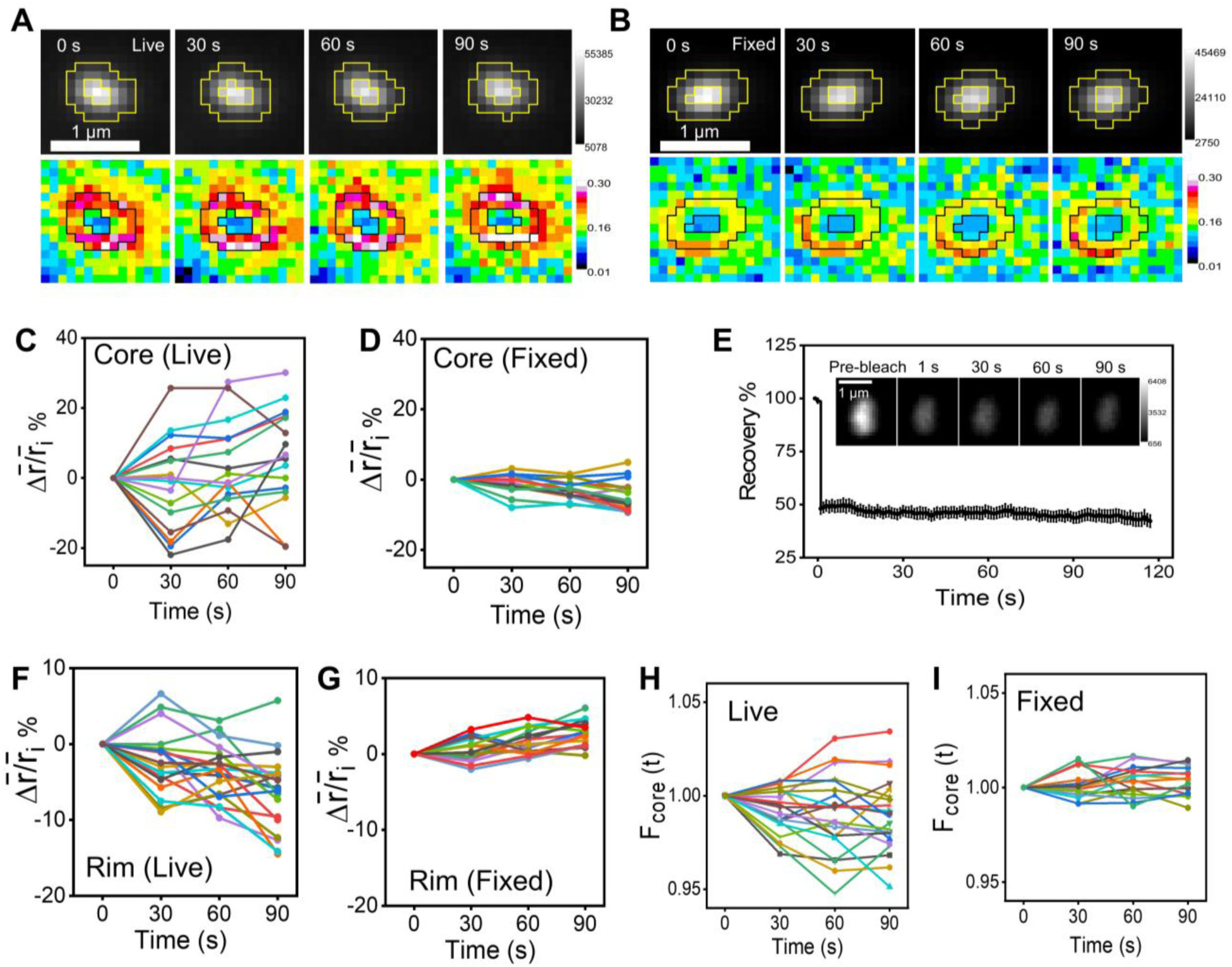
Intra-cluster spatiotemporal dynamics of TNFR1-EGFP reveal spontaneous rearrangements of receptor molecules within clusters. (**A** and **B**) Time-lapse intensity (top) and fluorescence anisotropy (bottom) maps of a single TNFR1-EGFP cluster in live (A) and fixed (B) HeLa cells monitored for 90 seconds. (**C** and **D**) Percentage change in relative mean fluorescence anisotropy (Δ 𝑟̅⁄𝑟̅_𝑖_), where Δ𝑟̅ = 𝑟̅(𝑡) − 𝑟̅_𝑖_, for the core region plotted as a function of time (𝑡) in live (C) and fixed (D) HeLa cells expressing TNFR1-EGFP, acquired over a 90-second time course. Here, 𝑟̅_𝑖_represents the initial mean anisotropy (𝑡 = 0), and 𝑟̅(𝑡) represents the mean anisotropy at time 𝑡. The data represent 2 cells each from two independent experiments. (**E**) Fluorescence recovery of TNFR1-EGFP clusters after photobleaching, monitored up to 120 seconds at 1-second intervals using confocal microscopy. Data are shown as mean ± SEM (n = 10 clusters from 2 cells). Representative confocal images of a TNFR1-EGFP cluster before and after photobleaching are shown in the inset. Images were processed to reduce noise by Gaussian averaging (radius = 1) using ImageJ. (**F** and **G**) Percentage change in relative mean fluorescence anisotropy (Δ 𝑟̅⁄𝑟̅_𝑖_) of the rim region plotted as a function of time (𝑡) in live (G) and fixed (H) HeLa cells expressing TNFR1-EGFP, acquired over a 90-second time course. (**H** and **I**) Temporal changes in the fractional fluorescence intensity of the core region, 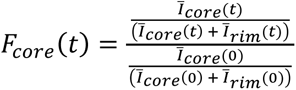, for TNFR1-EGFP clusters in live (H) and fixed (I) HeLa cells, acquired over a 90-second time course. Here, 𝐼_𝑐̅𝑜𝑟𝑒_(𝑡) and 𝐼_𝑟̅𝑖𝑚_(𝑡) represent mean fluorescence intensities in the core and rim regions at time 𝑡. Data in (C, D, F, G, H, I) represent n = 20 clusters from 2 cells each across two independent experiments. Scale bars are indicated in the corresponding images.

Since the intra-cluster reorganization of receptors may also alter the rim’s IRD, we analyzed whether our method can capture the associated temporal variations of homo-FRET in the rim. Comparison of the temporal variation of relative changes in 𝑟̅_𝑟𝑖𝑚_in live (**Fig. 2F**) to the clusters in fixed (**Fig. 2G**) cells demonstrated that our experimental arrangement can monitor the changes in the rim’s IRD. Intra-cluster reorganization of receptors in the cluster should lead to fluorescence redistribution across the cluster’s core and the rim. The higher fluctuations of normalized fractional intensity in the core (𝐹_𝑐𝑜𝑟𝑒_(𝑡)) in live (**Fig. 2H**) than in fixed (**Fig. 2I**) cells, reconfirmed intra-cluster reorganization of the TNFR1-EGFP.

### sTNF-α binding and PLAD inhibition differentially remodel TNFR1 IRD and signaling

The continuous intra-cluster reorganization observed in TNFR1-EGFP clusters (**Fig. 2**) prompted us to test whether sTNF-α binding enforces receptor rearrangement. We therefore examined changes in anisotropy maps following stimulation with sTNF-α or inhibition with zafirlukast (*34, 47*). Consistent with its role in activating TNFR1, sTNF-α increased nuclear NF-κB (p65) levels (*48*) with 20 ng/mL sufficient to induce maximal p65 activation within 15 minutes (**Fig. S4A**). For reproducibility, nuclear translocation was quantified at 30 minutes. Within 5 minutes, sTNF-α reduced TNFR1-EGFP cluster anisotropy in the core while increasing anisotropy in the rim (**Fig. 3, A and B**), indicating elevated IRD in the core and reduced IRD in the rim. This was accompanied by a modest but significant increase in cluster area within 30 minutes (**fig. S4B**), consistent with recruitment of additional receptors. These results support three hypotheses: (i) TNFR1 clusters contain both ligand-bound (**fig. S4C**) and ligand-free receptors, (ii) receptor rearrangement occurs on the experimental timescale, and (iii) ligand-bound receptors preferentially localize to the cluster core. Consequently, due to the higher tendency of ligand-bound TNFR1s to reside in the core, rim’s IRD decreased upon sTNF-α addition.

**Fig. 3.**
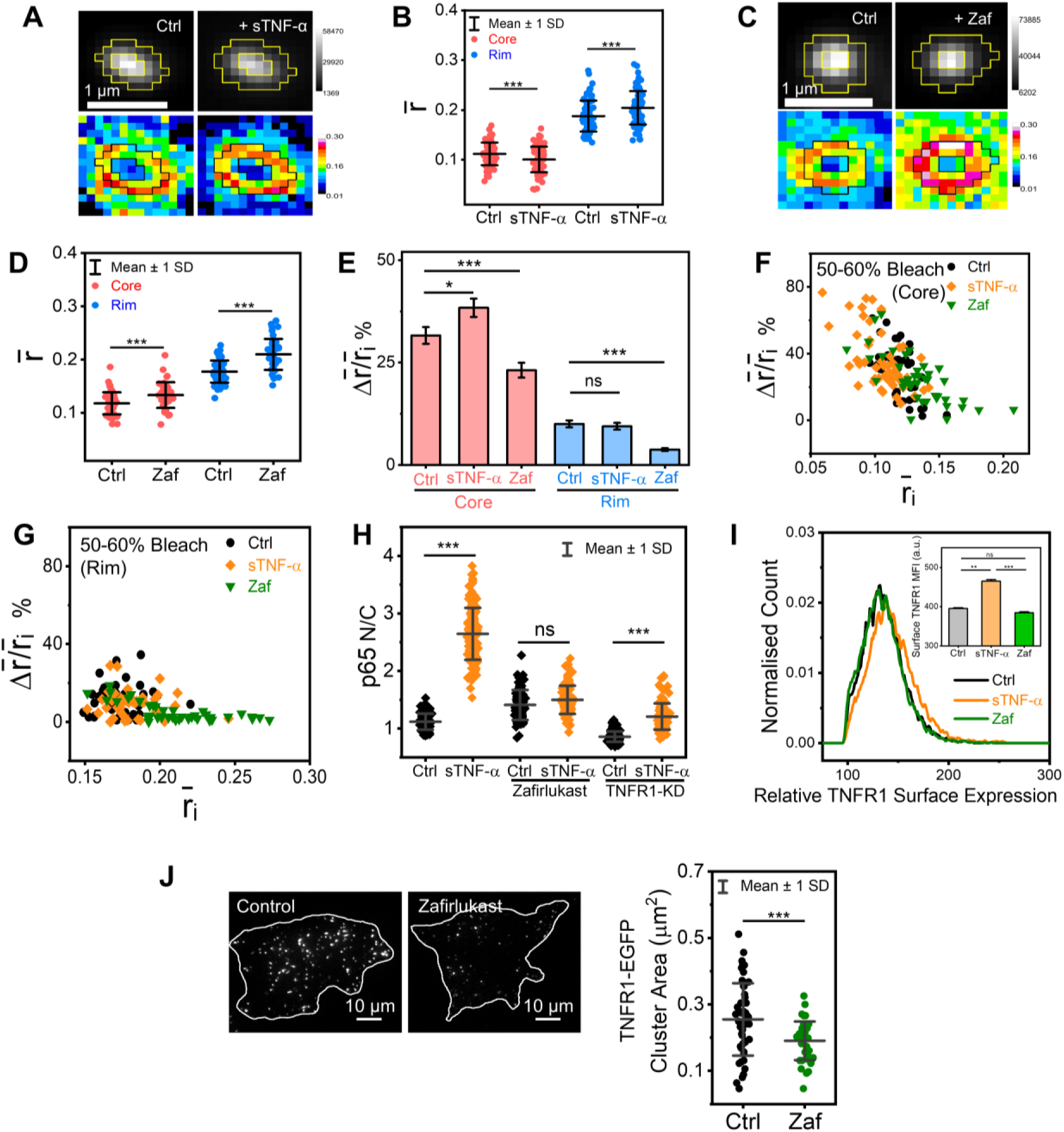
Ligand binding and PLAD inhibition differentially remodel TNFR1 IRD and signaling. (**A**) Representative fluorescence intensity (top) and corresponding fluorescence anisotropy (bottom) maps of the same TNFR1-EGFP cluster in live HeLa cells before (Ctrl) and after sTNF-α stimulation (20 ng/mL, 5 min). (**B**) Mean fluorescence anisotropy (𝑟̅) of the core (red) and rim (blue) regions of the same TNFR1-EGFP clusters (n = 90) under control (Ctrl) and sTNF-α–treated conditions. The data represent 3 cells each from three independent experiments. (**C**) Representative fluorescence intensity (top) and corresponding fluorescence anisotropy (bottom) maps of a TNFR1-EGFP cluster in live HeLa cells before (Ctrl) and after zafirlukast (Zaf) treatment (100 µM, 1 h). (**D**) Mean fluorescence anisotropy (𝑟̅) of the core (red) and rim (blue) regions of TNFR1-EGFP clusters (n = 50) under control (Ctrl) and zafirlukast-treated (Zaf) conditions. The data represent 3 cells each in three independent experiments. (**E**) Percentage change in relative mean fluorescence anisotropy (Δ 𝑟̅⁄𝑟̅_𝑖_) of the core (red) and rim (blue) regions of TNFR1-EGFP clusters (n ≥ 45) upon 50-60% photobleaching in control (Ctrl), sTNF-α, and zafirlukast-treated (Zaf) conditions. Here, Δ𝑟̅ = 𝑟_𝑓̅_ − 𝑟̅_𝑖_ where 𝑟_𝑓̅_ and 𝑟̅_𝑖_ represent post- and pre-bleach mean anisotropy, respectively. The data represent 2 cells each in three independent experiments. (**F** and **G**) Scatter plots showing the relationship between percentage change in relative mean fluorescence anisotropy (Δ 𝑟̅⁄𝑟̅_𝑖_) and the initial mean anisotropy (𝑟̅_𝑖_) in the core (F) and rim (G) regions of TNFR1-EGFP clusters (n ≥ 45) upon 50-60% photobleaching under control (Ctrl), sTNF-α, and zafirlukast-treated (Zaf) conditions. The data represent 2 cells each in three independent experiments. (**H**) Quantification of nuclear-to-cytoplasmic (N/C) mean intensity ratio of endogenous NF-κB (p65) in control (Ctrl) and sTNF-α– stimulated (20 ng/mL, 30 min) HeLa cells under untreated, zafirlukast-treated (Zaf), and TNFR1-knockdown (TNFR1-KD) conditions. Each control group corresponds to its respective treatment. The data represent 35 cells per condition from three independent experiments. (**I**) Flow cytometric analysis of surface TNFR1 expression in control (Ctrl), sTNF-α (20 ng/mL, 30 min), and zafirlukast-treated (Zaf) HeLa cells. The quantification of the median fluorescence intensity (MFI) is shown in the inset. (**J**) Representative TIRF images of immobile TNFR1-EGFP clusters in untreated (Ctrl, left) and zafirlukast-treated (Zaf, right) HeLa cells, with quantification of cluster area (μm²) shown (Ctrl: n = 49 clusters; Zaf: n = 43 clusters). Data are presented as means ± SD from 2 cells each in three independent experiments. Statistical significance between datasets was determined by paired two-tailed Student’s *t*-test (B), unpaired two-tailed Student’s *t*-test (D and J), or one-way ANOVA (E, H, I) and is indicated in the figures as follows: ∗P < 0.05; ∗∗P < 0.01; ∗∗∗P < 0.001; ns, not significant. Scale bars are indicated in the corresponding images.

Zafirlukast, a leukotriene receptor antagonist (*49*) that binds to the PLAD domain inhibiting TNFR1 PLAD-PLAD interaction, therefore it is expected to disrupt secondary clustering, effectively inhibiting sTNF–α–mediated NF-κB activation (*33, 34, 47*) at 100 μM within 1 hour (**Fig. S4D**). Unlike sTNF-α, zafirlukast caused a global reduction in IRD across both core and rim (**Fig. 3, C and D**), reflected by increased anisotropy values, while preserving cluster integrity. Photobleaching analysis showed that sTNF-α increased homo-FRET in the cluster core. In contrast, zafirlukast markedly reduced homo-FRET in both core and rim, as indicated by changes in relative mean anisotropy after 50–60% photobleaching (**Fig. 3E**). A negative correlation between initial anisotropy (𝑟̅_𝑖_) and photobleaching-induced anisotropy changes (**Fig. 3F**) established substantial inter-cluster IRD variation in the core. While sTNF-α minimally affected the rim, zafirlukast reduced IRD in both compartments (**Fig. 3G**), consistent with preferential localization of ligand-bound receptors in the core.

We next examined the relationship between IRD and signaling efficacy. Depletion of TNFR1 decreased the basal nuclear-to-cytoplasmic ratio of endogenous NF-κB (**fig. S4E**), indicating that basal signaling depends on receptor abundance (**Fig. 1H**). Conversely, TNFR1-EGFP overexpression **(Fig. 1G**) elevated basal NF-κB activity (**fig. S5A**), likely through increased surface receptor levels. For a given dose of sTNF-α, the increase of N/C ratio of NF-κB (**Fig. 3H and fig. S4E**), normalized to surface receptor levels in the cell, can be used as a quantitative indicator of ligand signaling efficacy. The effect of sTNF-α significantly diminished in TNFR1-depleted HeLa cells, confirming that p65 nuclear translocation is mediated through TNFR1 signaling. Zafirlukast markedly reduced sTNF-α efficacy (**Fig. 3H, and figs. S4E and S5B**) without altering surface or total TNFR1 abundance (**Fig. 3I and fig. S5C**). A slight increase in surface TNFR1 levels after 30 minutes of sTNF-α stimulation (**Fig. 3I and fig. S5D**) did not rescue signaling in the presence of zafirlukast (**Fig. 3H**). Thus, reduced signaling efficacy under zafirlukast reflects impaired IRD (**Fig. 3D**) rather than altered receptor number. Notably, NF-κB activation correlated with IRD changes-increased core IRD promoted signaling, whereas decreased IRD impeded activation (**Fig. 3, B, D, and H**). Finally, we assessed whether zafirlukast dislodged receptors from the clusters. TIRF imaging revealed a modest but significant reduction in TNFR1-EGFP cluster area (**Fig. 3J**), suggesting receptor loss, yet clusters were not fully disassembled, indicating that interactions beyond PLAD–PLAD contacts maintain structural integrity (*21*).

### Membrane tension affects the TNFR1 signaling by modulating IRD

Based on our finding that the IRD of TNFR1 clusters determines their signaling state (**Figs. 2 and 3**), we next examined how membrane tension, a factor known to modulate local curvature (*50–52*) and lipid packing (*53*) influences IRD and signaling. Consequently, HeLa cells were subjected to osmotic perturbations while monitoring membrane tension with Flipper-TR (*54*). Within 2 minutes of hypotonic treatment (75% water), cells displayed elevated membrane tension (**Fig. 4A and fig. S6A**), whereas hypertonic treatment reduced membrane tension after 2 minutes (150 mM mannitol; **Fig. 4B and fig. S6B**). These changes in tension directly altered cluster IRD: hypotonic stress increased fluorescence anisotropy in both core and rim, indicating reduced IRD (**Fig. 4C**), while hypertonic stress decreased anisotropy, reflecting an increase in IRD (**Fig. 4D**). We exploited membrane tension-induced changes in IRD (**Fig. 4, C and D**) to explore the relationship between receptor density and signaling efficiency (**Fig. 4E and fig. S6C**). The decrease in IRD under hypotonic stress resembled that seen with zafirlukast treatment (**fig. S6D**). Functionally, reduced IRD (**Fig. 4C**) did not independently trigger NF-κB translocation (**Fig. 4E**), whereas elevated IRD under hypertonic stress induced p65 translocation, even if less efficiently than sTNF-α stimulation (**Fig. 4E**). Western blot analysis of phosphorylated p65 further confirmed that sTNF-α failed to activate p65 under hypotonic conditions (**fig. S6E**). Importantly, NF-κB translocation was abolished in TNFR1-depleted cells (**Fig. 4E**), demonstrating that hypertonicity-induced activation of NF-κB is TNFR1-dependent. As no external sTNF-α was supplied, we conclude that TNFR1 activation occurred independently of sTNF-α stimulation.

**Fig. 4.**
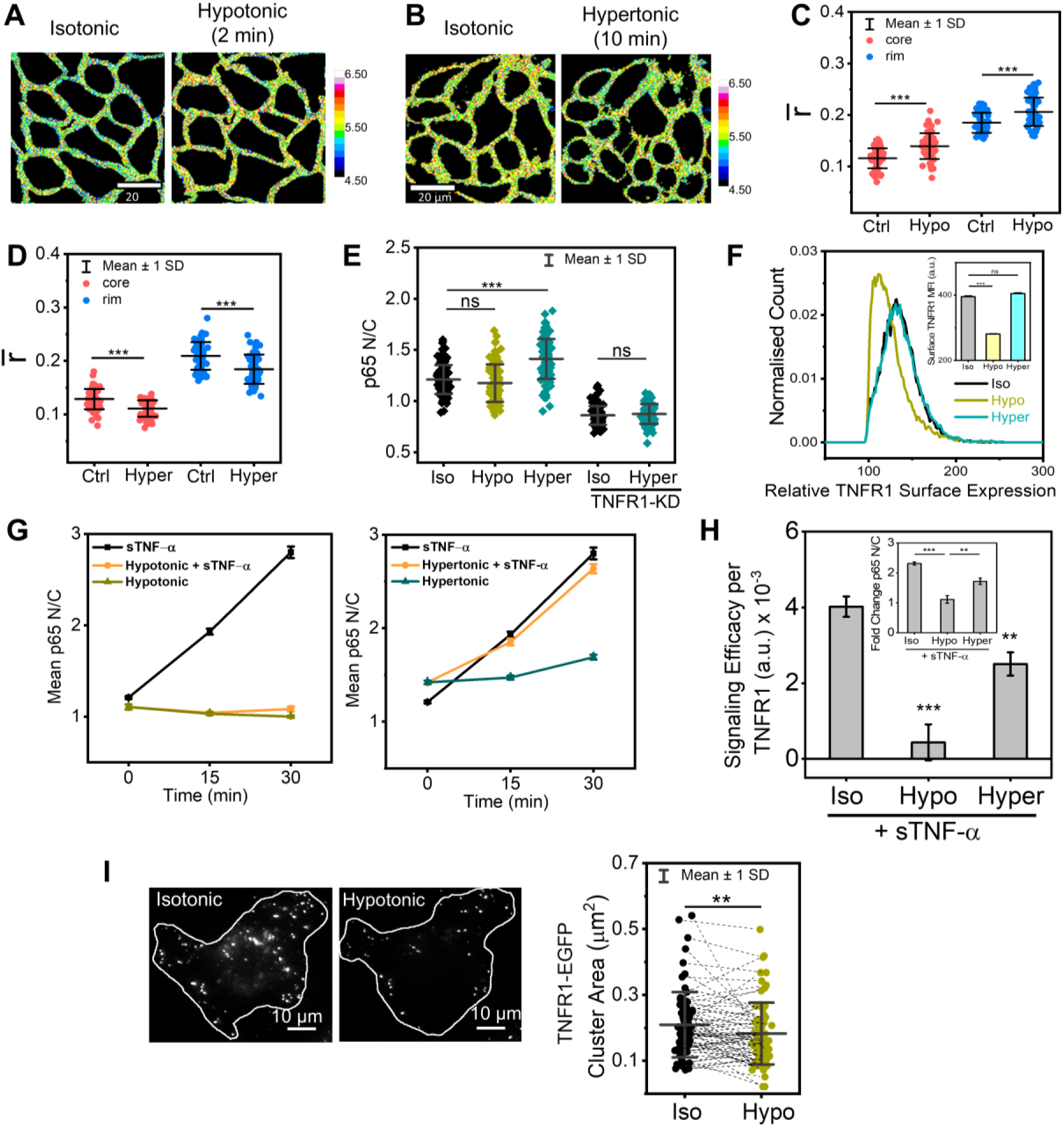
Membrane tension regulates TNFR1 IRD and modulates signaling efficacy. (**A** and **B**) Representative fluorescence lifetime maps of Flipper-TR (200 nM, 15 min) in HeLa cells subjected to (A) hypotonic (75% water, 2 min) and (B) hypertonic (150 mM mannitol, 10 min) shocks relative to the isotonic condition. (**C**) Mean fluorescence anisotropy (𝑟̅) of the core (red) and rim (blue) regions of TNFR1-EGFP clusters (n = 60) under hypotonic conditions (Hypo) compared to isotonic control (Ctrl). The data represent 5 cells each in three independent experiments. (**D**) Mean fluorescence anisotropy (𝑟̅) of the core (red) and rim (blue) regions of TNFR1-EGFP clusters (n = 60) under hypertonic conditions (Hyper) compared to isotonic control (Ctrl). The data represent 5 cells each in three independent experiments. (**E**) Nuclear-to-cytoplasmic (N/C) mean intensity ratio of endogenous NF-κB (p65) analyzed by immunostaining in HeLa cells subjected to isotonic (Iso), hypotonic (Hypo), or hypertonic (Hyper) conditions, in both control and TNFR1-knockdown (TNFR1-KD) cells. The data represent 35 cells each in three independent experiments. (**F**) Surface expression levels of endogenous TNFR1 in HeLa cells under isotonic (Iso), hypotonic (Hypo), and hypertonic (Hyper) conditions measured by flow cytometry. The quantification of the median fluorescence intensity (MFI) in each condition is shown in the inset. (**G**) Quantification of p65 N/C mean intensity ratio over 30 minutes following sTNF-α stimulation under hypotonic (left) and hypertonic (right) conditions compared to isotonic control. (**H**) Signaling efficacy of TNFR1 in HeLa cells subjected to isotonic (Iso), hypotonic (Hypo), and hypertonic (Hyper) shocks in response to sTNF-α. Signaling efficacy represents the N/C ratio change of p65 normalized to surface TNFR1 levels (ΔN/C p65 ratio / respective surface TNFR1 MFI). The inset shows fold change in p65 N/C ratio under each condition relative to corresponding unstimulated controls. Data represent means ± SD of 30 cells each in three independent experiments. (**I**) Representative TIRF images of TNFR1-EGFP–expressing HeLa cells before (left) and after (right) hypotonic shock. Quantification of TNFR1-EGFP cluster area (μm²) of the same clusters (n = 73) before and after treatment is shown. Data represent means ± SD from 3 cells each in three independent experiments. Statistical significance between datasets was determined by unpaired two-tailed Student’s *t*-test (C, D), one-way ANOVA (E, F, H), or paired two-tailed Student’s *t*-test (I) and is indicated in the figures as follows: ∗P < 0.05; ∗∗P < 0.01; ∗∗∗P < 0.001; ns, not significant. Scale bars are indicated in the corresponding images.

We next compared the response to exogenous sTNF-α under altered tension. While surface TNFR1 was reduced under hypotonic conditions, hypertonicity caused negligible changes (**Fig. 4F**), and total cellular TNFR1 levels remained unchanged across conditions **(fig. S5C**). Strikingly, sTNF-α failed to induce NF-κB nuclear translocation in hypotonic conditions **(Fig. 4G and fig. S6E**), consistent with suppressed signaling under reduced IRD. In contrast, hypertonic stress not only promoted ligand-free TNFR1 activation but also enhanced the response to sTNF-α, as confirmed by increased p65 phosphorylation (**Fig. 4G and fig. S6E**). Quantification revealed that signaling efficacy dropped sharply under hypotonic conditions, but only modestly under hypertonic conditions (**Fig. 4H).**

Finally, we examined the morphological changes in clusters using TIRF imaging. Hypotonic stress caused a small but significant decrease in TNFR1-EGFP cluster (**Fig. 4I and fig. S6F**), suggesting receptor loss from clusters. In contrast, hypertonicity had minimal effect on cluster area (**fig. S6G**). Together, these results indicate that increased membrane tension reduces cluster IRD and partial receptor dissociation, thereby impairing TNFR1 signaling, whereas decreased tension increases IRD and supports ligand-free receptor activation.

### Mechanical cues from a 3D gel-like microenvironment trigger TNFR1 signaling independent of ligand

Our previous work showed that transitioning THP-1 monocytes from a fluid-like 2D to a gel-like 3D extracellular microenvironment induces ligand-free NF-κB activation and differentiation into macrophages (*55*). To test whether TNFR1 mediates this response, we embedded HeLa cells in a 3D gel-like matrix. Similar to THP-1 cells, HeLa cells exhibited sTNF-α–independent nuclear translocation of NF-κB, as observed both with p65-EGFP (**fig. S7, A and B**) and by immunostaining of endogenous p65 (**Fig. 5A and fig. S7C**). TNFR1 inhibition (**fig. S7, A, B and C**) or depletion (**Fig. 5A**) abolished nuclear translocation of endogenous NF-κB and p65-EGFP in a 3D gel-like extracellular microenvironment, confirming that microenvironment-mediated NF-κB activation requires TNFR1. Notably, this activation occurred without changes in TNFR1 surface levels (**fig. S7D**).

**Fig. 5.**
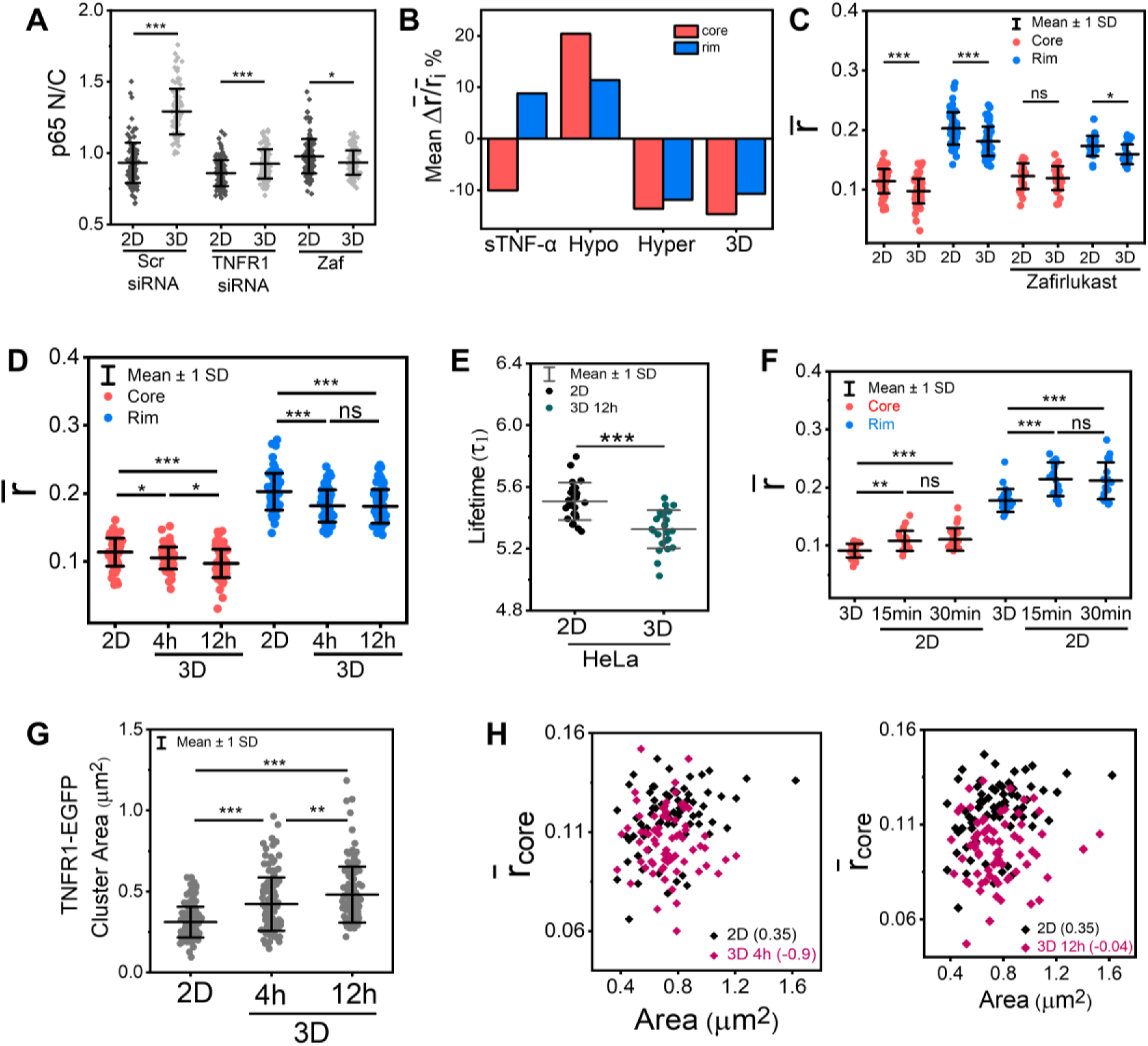
3D gel-like microenvironment induces ligand-independent TNFR1 signaling by modulating IRD. (**A**) Nuclear-to-cytoplasmic (N/C) mean intensity ratio of endogenous NF-κB (p65) in HeLa cells cultured in 2D (control) or 3D (1% agarose gel cover, 12 h) under scrambled siRNA (Scr siRNA), TNFR1 siRNA, or zafirlukast-treated (Zaf) conditions, analyzed by immunostaining. The data represent 35 cells each in three independent experiments. (**B**) Percentage change in relative mean fluorescence anisotropy (Δ 𝑟̅⁄𝑟̅_𝑖_) of core (red) and rim (blue) regions of TNFR1-EGFP clusters (n = 60) in HeLa cells under sTNF-α treatment, hypotonic (Hypo), hypertonic (Hyper), and 3D conditions, normalized to their respective controls. Here, Δ𝑟̅ = 𝑟_𝑓̅_ − 𝑟̅_𝑖_ where 𝑟_𝑓̅_ and 𝑟̅_𝑖_ represent the final and initial mean anisotropy for each treatment, respectively. The data represent 3 cells each in three independent experiments. (**C**) Mean fluorescence anisotropy (𝑟̅) of core (red) and rim (blue) regions of TNFR1-EGFP clusters in HeLa cells in 2D (control) or 3D under untreated (n = 60 clusters) or zafirlukast-treated (n = 25 clusters) conditions. The data represent 3 cells each in three independent experiments. (**D**) Mean fluorescence anisotropy (𝑟̅) of core (red) and rim (blue) regions of TNFR1-EGFP clusters (n = 70) during the transition from 2D to 3D culture after 4 h and 12 h of incubation. The data represent 3 cells each in three independent experiments. (**E**) Fluorescence lifetime (𝜏_1_) of Flipper-TR in HeLa cells cultured in 2D or 3D medium. The data represent 8 cells each in three independent experiments. (**F**) Mean fluorescence anisotropy (𝑟̅) of core (red) and rim (blue) regions of TNFR1-EGFP clusters (n = 20) in HeLa cells transitioning from 3D to 2D culture after 15 min and 30 min of incubation. The data represent 2 cells each in two independent experiments. (**G**) Quantification of area (μm²) of TNFR1-EGFP clusters (n = 124) in HeLa cells in 2D (control) and 3D conditions after 4 h and 12 h of incubation, acquired by epifluorescence imaging. The data represent 5 cells each in three independent experiments. (**H**) Scatter plots of TNFR1-EGFP cluster area (μm²) versus mean fluorescence anisotropy of the core (𝑟_𝑐̅𝑜𝑟𝑒_) of the same TNFR1-EGFP clusters (n = 70) in HeLa cells cultured in 2D and 3D [4 h (left) and 12 h (right)]. Pearson’s correlation coefficients are indicated for each condition. The data represent 3 cells each in three independent experiments. Statistical significance between datasets was determined by one-way ANOVA (A, D, F, G) or unpaired two-tailed Student’s *t*-test (C, E) and is indicated in the figures as follows: ∗P < 0.05; ∗∗P < 0.01; ∗∗∗P < 0.001; ns, not significant. Scale bars are indicated in the corresponding images.

**Fig. 6.**
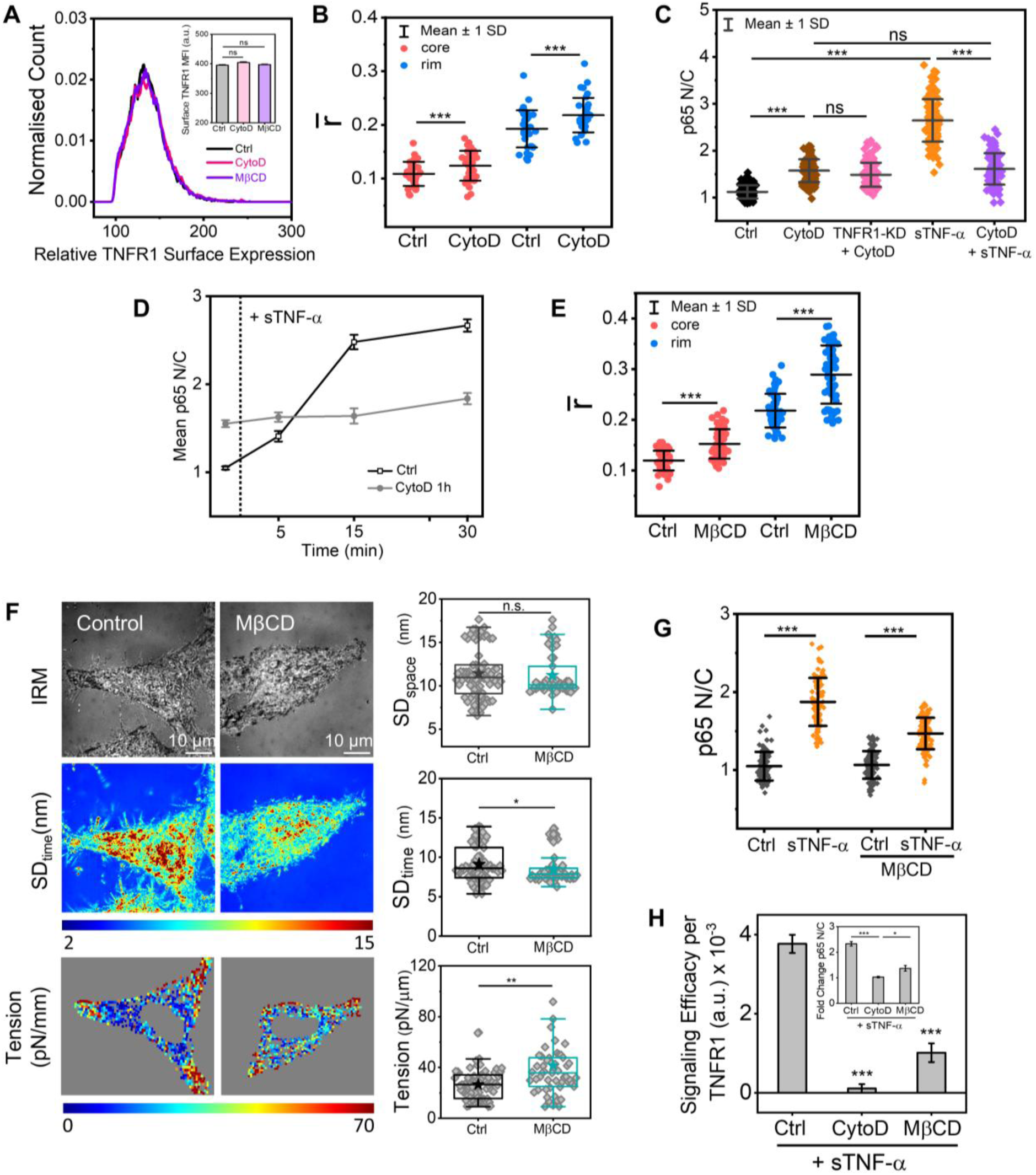
Cytoskeletal and membrane perturbations reduce TNFR1 IRD and signaling efficacy. (**A**) Surface expression of endogenous TNFR1 in HeLa cells under control (Ctrl), cytochalasin D-treated (CytoD, 2 μM, 1 h), and methyl-β-cyclodextrin-treated (MβCD, 2 mM, 1 h) conditions, measured by flow cytometry. Median fluorescence intensity (MFI) is shown in the inset. (**B**) Mean fluorescence anisotropy (𝑟̅) of the core (red) and rim (blue) regions of TNFR1-EGFP clusters (n = 34) in control (Ctrl) and CytoD-treated HeLa cells. The data represent 2 cells across two independent experiments. (**C**) Nuclear-to-cytoplasmic (N/C) mean intensity ratio of NF-κB (p65) in control (Ctrl), CytoD-treated, TNFR1-knockdown (TNFR1-KD), and sTNF-α-stimulated HeLa cells, analyzed by immunostaining. The data represent 35 cells each in three independent experiments. (**D**) Temporal changes in the N/C ratio of p65 in control (Ctrl) and CytoD-treated HeLa cells in response to sTNF-α, monitored for 30 minutes. Data are presented as means ± SEM of 35 cells in three independent experiments. (**E**) Mean fluorescence anisotropy (𝑟̅) of the core (red) and rim (blue) regions of TNFR1-EGFP clusters (n = 64) in control (Ctrl) and MβCD-treated HeLa cells. The data represent 3 cells across three independent experiments. (**F**) Representative Interference Reflection Microscopy (IRM) images (top), corresponding SD_time_ maps (middle), and membrane tension maps (bottom) of control (Ctrl) and MβCD-treated HeLa cells. Quantification of SD_time_ (nm) and membrane tension (pN/μm) is shown (Ctrl: n = 79 cells; MβCD: n = 57 cells). Data are representative of three independent experiments. (**G**) N/C mean intensity ratio of NF-κB (p65) in control (Ctrl) and MβCD-treated HeLa cells with or without sTNF-α stimulation. The data represent 35 cells each in three independent experiments. (**H**) TNFR1 signaling efficacy, defined as the change in p65 N/C ratio normalized to surface TNFR1 levels (MFI), in control, CytoD, and MβCD-treated HeLa cells in response to sTNF-α. The fold change of p65 N/C, normalized to its respective basal levels, is shown in the inset. Data are presented as means ± SD of 30 cells each in three independent experiments. Statistical significance between datasets was determined by one-way ANOVA (A, C, G, H) or unpaired two-tailed Student’s *t*-test (B, E) and is indicated in the figures as follows: ∗P < 0.05; ∗∗P < 0.01; ∗∗∗P < 0.001; ns, not significant. Scale bars are indicated in the corresponding images.

**Fig. 7.**
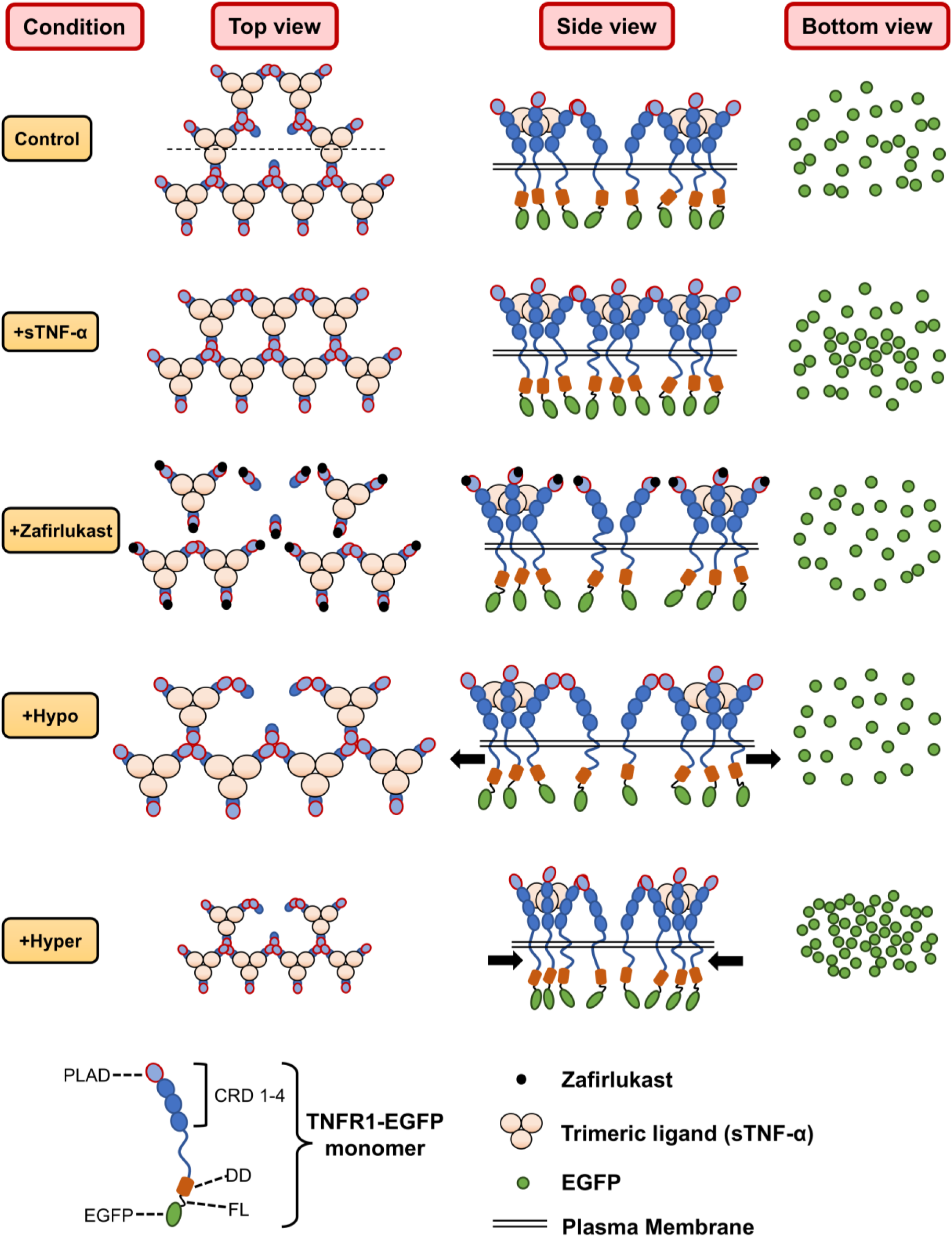
Schematic model of TNFR1 cluster organization governed by ligand-receptor and receptor-receptor interactions. Top (extracellular), side (lateral), and bottom (cytoplasmic) views illustrate the proposed schematic arrangement of TNFR1–EGFP molecules within a cluster under different conditions. In the native (control) state, TNFR1 exists in pre-assembled clusters formed through ligand–receptor and PLAD–PLAD interactions, with a few unoccupied binding sites in the cluster core. Upon ligand stimulation, these vacant sites become occupied, resulting in increased intermolecular proximity (higher IRD) in the core and reduced IRD at the rim, as visualized from the cytoplasmic (bottom) view. Zafirlukast treatment disrupts PLAD–PLAD interactions, leading to decreased IRD in both the core and rim regions. Hypotonic stress increases membrane tension, expanding intermolecular distances and reducing IRD throughout the cluster. Conversely, hypertonic stress reduces membrane tension, promoting molecular proximity and increasing IRD across both regions. PLAD, pre-ligand assembly domain; CRD, cysteine-rich domain; DD, death domain; FL, flexible linker.

We next examined how a 3D microenvironment influenced IRD. In contrast to sTNF-α, which increased IRD in the core and reduced it in the rim (**Fig. 3B**), or osmotic perturbations, which altered both regions in the same direction (**Fig. 5C**), the 3D gel-like condition increased IRD simultaneously in both the core and rim (**Fig. 5B**). This effect was blocked by zafirlukast, linking IRD changes to TNFR1 signaling (**Fig. 5C)**. Because same-directional IRD changes in core and rim are indicative of altered membrane tension, we hypothesized that a 3D matrix lowers membrane tension. Consistent with this, cells transitioning from 2D to 3D gradually exhibited higher IRD in both core and rim (**Fig. 5D**), while Flipper-TR lifetime measurements confirmed reduced membrane tension in both HeLa (**Fig. 5E**) and THP-1 cells (**fig. S7E**). Reversing cells from 3D back to 2D restored IRD within 15 minutes (**Fig. 5F**), indicating tension-dependent reversibility of the IRD.

Structurally, the 3D microenvironment increased TNFR1-EGFP cluster area (**Fig. 5G),** but no correlation was found between cluster size and IRD **(Fig. 5H and fig. S7F).** Together, these findings suggest that 3D gel-like environments activate TNFR1 signaling in a ligand-independent manner by reducing membrane tension and increasing IRD across the cluster in addition to increasing its size.

### Structural integrity of actin and cholesterol is essential for TNFR1 signaling

Given the role of the cytoskeleton in membrane organization and receptor signaling, we next tested whether actin integrity affects TNFR1 clustering and signaling. Disruption of actin filaments with cytochalasin D (CytoD) did not alter surface TNFR1 levels (**Fig. 6A**) or total TNFR1 abundance (**fig. S5C**), but it reduced IRD in both the core and rim of clusters **(Fig. 6B**). Functionally, CytoD induced NF-κB translocation even in TNFR1-depleted cells (**Fig. 6C and fig. S8A)**, consistent with reports that CytoD activates NF-κB through ROS generation (*56*). Indeed, the ROS quencher vitamin C (**fig. S8B**) (*57–59*) abolished CytoD-induced NF-κB nuclear translocation (**fig. S8, C and D)** without affecting cluster IRD (**fig. S8E**). Notably, vitamin C had no impact on sTNF-α–induced NF-κB activation in untreated cells (**fig. S8, C and D**). When comparing signaling kinetics, CytoD-treated cells exhibited delayed NF-κB translocation in response to sTNF-α relative to control cells (**Fig. 6D**), consistent with reduced IRD impairing TNFR1 signaling efficiency.

We next investigated whether membrane cholesterol, another key determinant of receptor organization (*60*), influences TNFR1 clustering and signaling. Cholesterol depletion with methyl-β-cyclodextrin (MβCD) significantly reduced IRD in TNFR1 clusters (**Fig. 6E**), likely through elevated membrane tension (**Fig. 6F**) (*61*) and/or direct effects on clustering (*60*). Cholesterol depletion caused negligible changes in cluster area (**fig. S9A**) and did not affect surface TNFR1 levels (**Fig. 6A**), suggesting that receptor loss from clusters was minimal. Despite preserved receptor abundance, cholesterol depletion impaired sTNF-α–mediated NF-κB activation (**Fig. 6G and fig. S9B**). Together, these findings demonstrate that both actin disruption and cholesterol depletion reduce TNFR1 IRD without significantly altering receptor abundance, thereby diminishing the signaling efficacy of TNFR1 clusters in response to sTNF-α (**Fig. 6H**).

## Discussion

Clustering of transmembrane receptors is a conserved mechanism for initiating signaling across diverse systems, including receptor tyrosine kinases (*44, 62*), T-cell receptors (*63*), bacterial chemoreceptors (*64*), and TNFRs (*1, 14*). In the TNFR superfamily, ligand-bound TNF–TNFR trimers pre-assemble into higher-order clusters (*22*) where the extracellular PLAD domain organizes into a hexagonal lattice (**Fig. 7 and fig. S4C)** (*65, 66*).

On the cytoplasmic side of TNFR1, the death domain is connected to the membrane by a long, flexible 122–amino acid loop (>41 nm), followed by an additional 17–amino acid loop (5.8 nm) at the C-terminus (Uniprot ID: P19438). To visualize receptor organization, we inserted an extra linker (7–21 amino acids; 2.4-7.1 nm) between TNFR1 and the EGFP tag. The extended cytoplasmic loops together with the engineered flexible linkers are therefore expected to provide high conformational flexibility to the EGFP tag. Instead of aligning with the ordered extracellular PLAD lattice, the tags likely adopt variable positions and orientations, producing heterogeneous EGFP distributions within TNFR1 nanoclusters (**Fig. 1, D and I, and Fig. 7**).

In our study, the nuclear translocation of NF-κB served as an indicator of TNFR1 signal activation. Stimulation with sTNF-α induces rapid NF-κB activation, with p65 nuclear translocation observed within five minutes (**fig. S4A and Fig. 6D**), even though surface-level TNFR1s remain unchanged during this time (**fig. S5D**). Subtle increases in cluster size appear only after 30 minutes of stimulation (**fig. S4B**), suggesting that local receptor reorganization—rather than cluster enlargement—is the primary trigger for signaling. This is supported by the rapid increase in IRD detected within five minutes of sTNF-α stimulation (**Fig. 3, A and B**). Manipulating TNFR1 expression further supports the role of clustering in signaling output: overexpression of TNFR1– EGFP enlarges exogenous cluster size (**fig. 1J**) and enhances p65 nuclear translocation (**fig. S5A**), whereas TNFR1 knockdown reduces endogenous and exogenous cluster size and lowers basal NF-κB activity (**Fig. 1J, fig. S2D, and fig. S4E**). Surface levels of TNFR1-EGFP could not be directly measured by FACS because the anti-TNFR1 antibody detects both exogenous and endogenous receptors. Together, these results indicate that TNFR1 signaling is governed by both IRD and cluster size, with IRD serving as the more sensitive determinant.

To quantify IRD, we used homo-FRET as a readout of IRD by measuring steady-state fluorescence anisotropy of TNFR1-EGFP, which offers advantages for homo-oligomeric receptors by avoiding stoichiometric imbalances seen with hetero-FRET measurements (*67*). The FRET efficiency is strongly dependent on the proximity of the receptors (2-10 nm), therefore making homo-FRET a good indicator of IRD changes. A negative correlation between anisotropy and photobleaching-induced FRET recovery confirmed that anisotropy maps reflect IRD distribution (**Fig. 1F and Fig. 3F**). Importantly, IRD was largely unaffected by variations in TNFR1 expression (**Fig. 1I**), confirming that our measurements were not biased by overexpression artifacts and supporting the cluster size-dependent activation. Our data reveal that TNFR1 clusters are highly dynamic, with receptors undergoing spontaneous intra-cluster rearrangement (**Fig. 2**). We suggest that the spontaneous intra-cluster reorganization of TNFR1 clusters, together with available vacant ligand-binding sites, enables a signaling output that scales with the dose of soluble TNF-α (**fig. S4A**). Thus, ligand stimulation further reorganizes receptors within clusters to modulate IRD in the core (**Fig. 3, A and B**). Pharmacological reduction of IRD with zafirlukast (**Fig. 3, C and D**) decreased cluster size (**Fig. 3J**) without altering surface receptor levels (**Fig. 3I**), demonstrating that IRD modulation is sufficient to regulate TNFR1 signaling state (**Fig. 3H**). Physical perturbations of the membrane microenvironment also modulate IRD. Hypertonic, hypotonic, and 3D agarose environments, which alter membrane tension, significantly shifted IRD. To independently determine the changes associated with membrane tension, we used the mechanosensitive probe Flipper-TR, which reports on variations in lipid order packaging under hypotonic (**Fig. 4A and fig. S6A**), hypertonic (**Fig. 4B and fig. S6B**), and 3D environments (**Fig. 5E and fig. S7E**). Disruption of actin filaments with cytochalasin D reduced IRD in both the core and rim of TNFR1 clusters and delayed NF-κB translocation in response to sTNF-α. This indicates that actin integrity is important for maintaining optimal TNFR1 cluster organization and signaling output. Similarly, cholesterol depletion using MβCD decreased IRD and impaired sTNF-α-mediated NF-κB activation, without significantly altering receptor abundance or cluster area. These results suggest that membrane cholesterol contributes to the structural organization of TNFR1 clusters and thereby modulates signaling efficacy. Since MβCD directly alters lipid composition by depleting membrane cholesterol (*68*), we instead relied on interference reflection microscopy (IRM) to assess membrane tension under cholesterol depletion (**Fig. 6F)** (*69*). Increased IRD led to ligand-independent activation of TNFR1 signaling, whereas decreased IRD rendered clusters unresponsive to sTNF-α, indicating that IRD functions as a molecular switch controlling signaling competence. In a hypertonic condition, the increase in IRD is accompanied by a negligible change in cluster size (**fig. S6G**) and surface level of receptors (**Fig. 4F**), strongly suggesting ligand-independent and IRD-mediated TNFR1 activation. Interestingly, hypertonic stress induced p65 nuclear translocation without detectable Ser536 phosphorylation (**fig. S6E**), implying activation through alternative p65 modifications (*70*). We also note an apparent increase in fluorescence intensity and the number of visible clusters under hypertonic conditions (**fig. S6G**). However, given the short exposure period, such an effect is unlikely to reflect a genuine rise in receptor abundance. Flow cytometry confirmed unchanged surface TNFR1 levels (**Fig. 4F**). We attribute this apparent rise to an increase in the penetration depth of the TIRF evanescent field due to refractive index changes in the hypertonic medium and to the cell shrinkage that likely brings cytoplasmic clusters closer to the membrane, making them detectable in TIRF imaging (**fig. S6G**).

We also addressed potential methodological concerns regarding EGFP tagging and anisotropy measurements. The use of monomeric EGFP connected by long flexible loops and a linker minimized steric hindrance and orientation artifacts, and functional assays confirmed signaling competence (**fig. S5A**). In conclusion, we demonstrate that IRD is a critical determinant of TNFR1 signaling, capable of modulating receptor activity independently of ligand binding. Our findings highlight IRD as a biophysical parameter that integrates membrane environment, receptor expression, and cytoskeletal architecture to fine-tune TNFR1-mediated NF-κB activation. This approach provides a framework for studying clustering-dependent regulation of other receptors and may have implications for pathological conditions where receptor clustering is dysregulated.

## Materials and Methods

### Plasmids

The following plasmids were obtained from Addgene: pTNFR1-tdEOS, a contribution from Mike Heilemann (#98273); mEGFP-N1, provided by Michael Davidson (#54767); and EGFP-p65, gifted by Johannes A. Schmid (#111190). The 2GFP (GFP-GFP) plasmid was generously provided by Prof. Maria Vartiainen from the University of Helsinki, Finland. The TNFR1-EGFP plasmid was constructed by sub-cloning the TNFR1 gene from the pTNFR1-tdEOS plasmid into the N-terminal of the mEGFP-N1 plasmid, with a flexible linker of 14 amino acids in between. The full-length human TNFR1 gene sequence was amplified by polymerase chain reaction (PCR) using the following primers:

5’-cggagctc**gccacc**ATGGGCCTCTCCACCGTGCCTG-3’ (forward)

5’-gtgtcgactcTCTGAGAAGACTGGGCGCGGGCGGGAGGGC-3’ (reverse)

The overhangs containing the restriction endonuclease sites (underlined) and the Kozak sequence (bold) are indicated in lowercase. The purified PCR product was inserted into the multiple cloning site (MCS) of the mEGFP-N1 vector backbone to generate C-terminally tagged TNFR1-EGFP. SacI (QuickCutTM Sac I, Takara Bio, #1627) and SalI (QuickCutTM Sal I, Takara Bio, #1636) restriction endonucleases were used. The digested products were ligated using T4 DNA Ligase (New England Biolabs, #M0202S). The fidelity of the chimeric clone was confirmed through diagnostic restriction digestion and direct sequencing (Eurofins, Bangalore, India).

### Cell culture and transfection

HeLa cell line was obtained from the National Center for Cell Science (NCCS) in Pune, India, while THP-1 cell line of American Type Culture Collection (ATCC) origin was generously provided by Dr. Prosenjit Sen from the Indian Association for the Cultivation of Science in Kolkata, India. Both cell lines were cultured in Dulbecco’s Modified Eagle Medium (DMEM) (HiMedia, #AL007G) supplemented with 10% fetal bovine serum (FBS) (HiMedia, #RM9955) and 1% penicillin-streptomycin (Antibiotic Solution 100X Liquid, HiMedia, #A001) at 37°C in a humidified incubator with 5% CO2.

HeLa cells were transfected with the afore-mentioned plasmids using FuGENE® HD Transfection Reagent (Promega, #E2311), following the manufacturer’s protocol. Approximately 1 x 10^5^ cells were seeded in glass-bottom Petri dishes 24 hours before transfection and allowed to reach 70% confluency. Transfection was carried out using 1 μg of plasmid DNA under serum-starved conditions for the first six hours after transfection, and then the cells were incubated with added culture media overnight at 37°C with 5% CO_2_. The medium was then replaced with DMEM supplemented with FBS and antibiotics before proceeding with further experiments. For TNFR1-EGFP overexpression-dependent signaling and IRD study, HeLa cells were transfected with 0.1, 0.5, and 1 μg of TNFR1-14aa-EGFP plasmid DNA. Linker length-dependent IRD studies were carried out by transfecting HeLa cells with 1 μg of TNFR1-7aa-EGFP, TNFR1-14aa-EGFP, and TNFR1-21aa-EGFP plasmid DNA. To generate TNFR1 knockdown cells, HeLa cells were transfected with pre-designed siRNA targeting human TNFRSF1A (Sigma-Aldrich/Merck, #SASI_Hs01_00033456, #SASI_Hs01_00033458), along with MISSION® siRNA Universal Negative Control #1 (Sigma-Aldrich/Merck, #SIC001) as the corresponding scrambled control. The transfection was performed using 30, 60, 90, and 120 nM of siRNA and FuGENE® under serum-starved conditions following the standard protocol for the optimization of TNFR1 knockdown efficiency. The transfection mix was incubated at room temperature for 15 minutes before being added drop-wise to the cells. The silenced cells were analyzed 96 hours post-transfection. The optimized 30 nM human TNFRSF1A siRNA was used for the experiments related to knockdown, IRD, and signaling measurements, as described in the result sections. To validate TNFR1-mediated NF-κB nuclear translocation in a 3D gel-like microenvironment, cells were pre-treated with 50 μM zafirlukast (Sigma-Aldrich/Merck, #Z4152) for 16 hours before proceeding with EGFP-p65 transfection.

### sTNF-α, drug, and osmolyte treatments

To initiate cytokine stimulation, HeLa cells (1 x 10^5^) were plated in glass-bottomed Petri dishes and incubated overnight at 37°C with 5% CO_2_. Subsequently, to deplete secreted cytokines and minimize cellular metabolic activity, the existing DMEM was aspirated, and cells were subjected to serum starvation for 4 hours. Afterward, cells were thoroughly rinsed twice with serum-free DMEM to eliminate any soluble tumor necrosis factor-alpha (sTNF-α) that might be present in the medium. Recombinant human TNF-α protein (ABclonal, #RP00993) was then introduced to the cells at various concentrations (5-100 ng/mL) and for different durations (5-45 minutes), as specified in the results section.

To evaluate the inhibitory effect of Zafirlukast on TNFR1-mediated signaling in HeLa cells, the compound was tested at varying concentrations and incubation times, as described in the Results section. For all subsequent experiments, HeLa cells were subjected to serum starvation for 3 hours, followed by a 1-hour pre-incubation with 100 µM Zafirlukast (Sigma-Aldrich/Merck, #Z4152). To induce osmotic shock in HeLa cells, the standard isotonic DMEM was replaced with hypertonic media (150 mM mannitol (Sigma-Aldrich/Merck, #M9647) dissolved in DMEM) or hypotonic media (75% autoclaved distilled water mixed with 25% DMEM by volume) for the duration specified in the results section. Additionally, to disrupt the actin cytoskeleton, HeLa cells were treated with 2 μM cytochalasin D (CytoD) (Sigma-Aldrich/Merck, #C8273) under serum-starved conditions for 1 hour, unless otherwise stated. Cholesterol depletion in HeLa cells involved a serum starvation for 2 hours, followed by 1 hour incubation with 2 mM methyl-β-cyclodextrin (MβCD) (Sigma-Aldrich/Merck, #C4555). Endocytosis inhibition was performed in TNFR1-EGFP-overexpressing HeLa cells by treating the cells with 80 μM dynasore (Sigma-Aldrich/ Merck, #D7693) for 50 minutes under serum-starved conditions. Prior to dynasore treatment, cells were preincubated in serum-free medium for 1 hour.

### Immunofluorescence

HeLa cells were seeded in a glass-bottom plate and incubated overnight in cell culture conditions. They were then fixed with 4% paraformaldehyde (PFA) (HiMedia, #GRM3660) in 1x phosphate-buffered saline (PBS) for 15 minutes. For immunostaining of the plasma membrane-associated TNFR1, the fixed cells were blocked with a 5% BSA solution in 1x PBS for 1 hour at room temperature, without permeabilizing the plasma membrane. After three washes with 1x PBS, the cells were incubated with a 1:500 dilution of TNFR1/TNFRSF1A rabbit pAb (Invitrogen, # PA5-95585), in antibody dilution buffer (3% BSA in 1x PBS) overnight at 4°C. Subsequently, the cells were washed thrice with 1x PBS and incubated with Alexa Fluor-594 conjugated secondary antibody (1:200 dilution, Anti-rabbit IgG, Cell Signaling Technology, #8889) for 1 hour at room temperature. After three washes with 1x PBS, the cells were ready for imaging. To validate our TNFR1-EGFP chimeric construct, 18-20 hours post-transfected, HeLa TNFR1-EGFP cells were fixed and prepared for membrane-associated TNFR1 immunostaining following the protocol described above. This approach allowed us to visualize most plasma-membrane-associated TNFR1-EGFP expressions (**Fig. S1B**) of the TNFR1-EGFP chimeric construct within the cells.

For immunostaining of NF-κB (p65) after fixation of the HeLa cells with 4% PFA in 1x PBS for 15 mins, they were permeabilized with 0.1% Triton X-100 (Sisco Research Laboratories, #64518) in 1x PBS for 4-5 mins. Then the fixed cells were blocked with a 5% BSA solution in 1x PBS for 1 hour at room temperature. After three washes with 1x PBS, the cells were incubated with a 1:500 dilution of NF-κB p65 mouse mAb (Cell Signaling Technology, #6956) in antibody dilution buffer (3% BSA in 1x PBS) overnight at 4°C. Subsequently, the cells were washed thrice with 1x PBS and incubated with Alexa Fluor-594 conjugated secondary antibody (1:200 dilution, Anti-mouse IgG, Cell Signaling Technology, #8890) for 1 hour at room temperature. After three washes with 1x PBS, the cells were ready for imaging. The nuclei were counterstained with Hoechst 33342 (ThermoFisher, #62249) (0.5 µg/mL, 5 minutes).

### Generation of TNFR1–EGFP Constructs with Altered Linker Lengths

To examine the effect of linker flexibility on TNFR1 organization and signaling, TNFR1–EGFP fusion constructs were generated containing short (7 amino acids) and long (21 amino acids) linkers, derived from the parental construct with a 14–amino acid flexible linker (TNFR1-14aa-EGFP). The cloning was performed by Gibson assembly (GeneArt™ Gibson Assembly® HiFi Cloning Kit, Invitrogen, # A46627) using primers designed to modify the linker region between TNFR1 and mEGFP.

For the short-linker construct (TNFR1-7aa-EGFP), the entire plasmid was PCR-amplified using Phusion™ High–Fidelity DNA Polymerase (ThermoFisher, # F530S) with primers that deleted 21 base pairs from the original linker sequence, thereby halving its length while maintaining the reading frame. Primer sequences were as follows (lowercase letters denote Gibson overhangs):

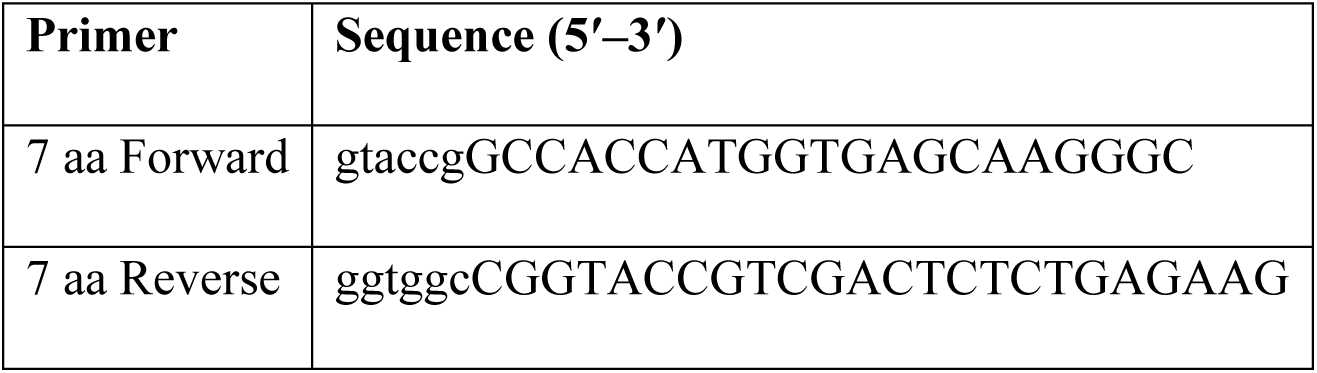

Following Gibson assembly, recombinant plasmids were electroporated into *E. coli* strain MC1061. Colonies were screened by BamHI digestion (QuickCut™ BamHI, Takara Bio, #1605), as the shortened linker design eliminated a BamHI site, enabling rapid verification.

For the long-linker construct (TNFR1-21aa-EGFP), PCR amplification was performed with primers that inserted 21 additional base pairs encoding a glycine–serine (GS)-rich motif (GGSGSSG) into the linker region, extending it to 21 amino acids. This flexible GS sequence was selected to preserve structural integrity and prevent steric interference (*71*). Primer sequences were:

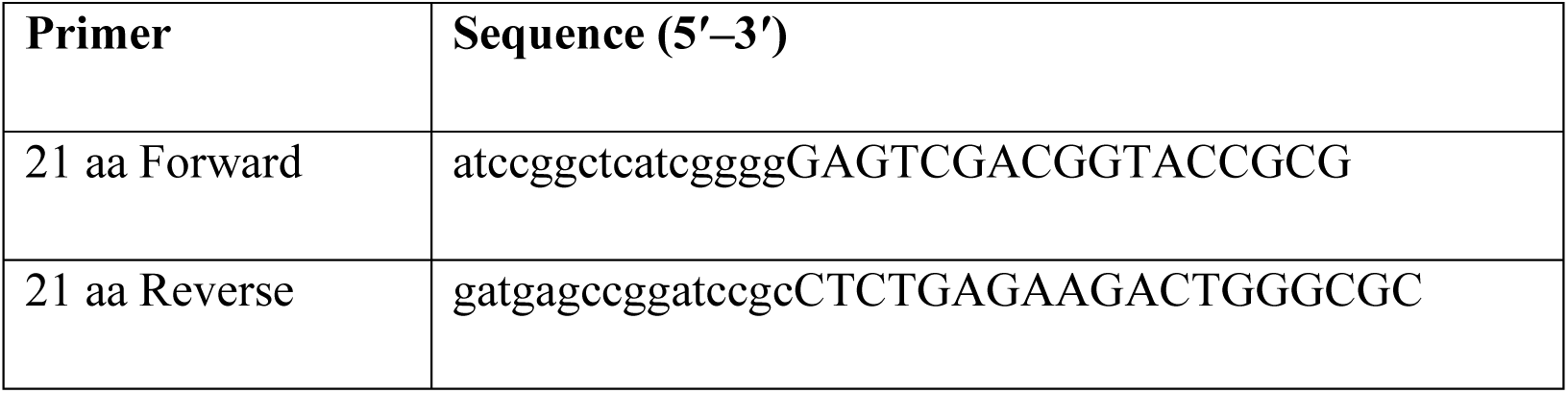

Following Gibson assembly, plasmids were electroporated into *E. coli* MC1061, and transformants were screened by PCR using a promoter-specific forward primer and a reverse primer annealing to the newly inserted linker sequence. Amplification was observed only from the modified construct and not from the parental TNFR1–EGFP plasmid, confirming the successful generation of the TNFR1–21aa–EGFP plasmid. Verification primers were:

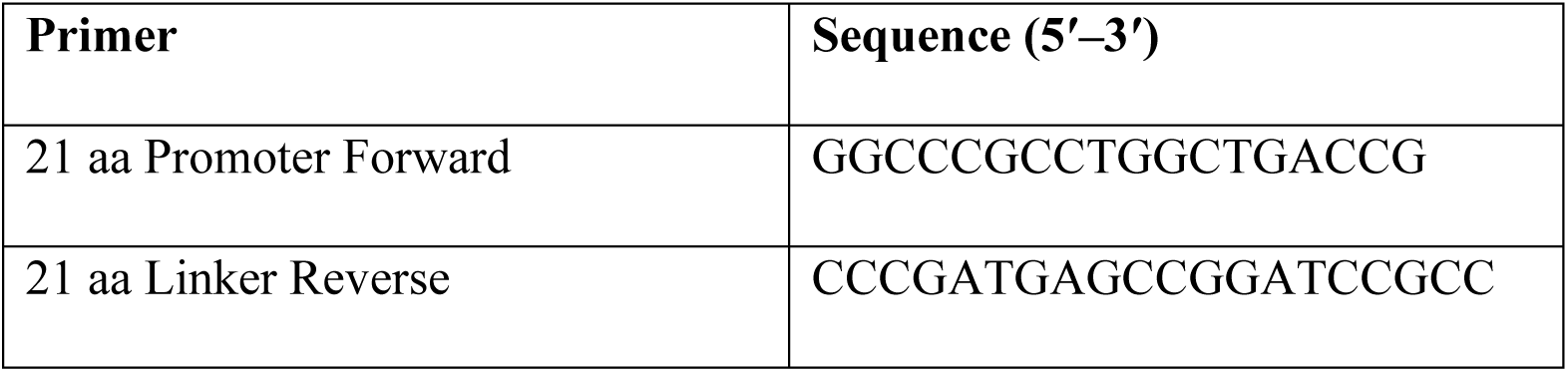

All constructs were validated by Sanger sequencing (Eurofins, Bangalore, India) to confirm the fidelity of the linker modifications.

### Cell extracts and immunoblotting

HeLa cells (1 x 10^5^) were cultured in 35 mm dishes and incubated overnight at 37°C with 5% CO2 until reaching approximately 70% confluency. Following this, the cells were either transfected with siRNA or treated with various drugs for specified durations before proceeding with cell lysate preparation. The cells were rinsed with ice-cold PBS and then lysed using 100 μL of RIPA buffer (50 mM Tris HCl, 150 mM NaCl, 1.0% (v/v) NP-40, 0.5% (w/v) sodium deoxycholate, 1.0 mM EDTA, 0.1% (w/v) SDS, and 0.01% (w/v) sodium azide, pH 7.4), supplemented with Protease and Phosphatase Inhibitor Cocktail (Sigma-Aldrich/Merck, #PPC1010). The lysates were incubated on ice for 15 minutes, and then the contents of each dish were transferred to the respective 1.5 mL microcentrifuge tubes. Total cell extracts were obtained by agitating the lysates at 4°C for 30 minutes. The cell lysates were centrifuged at 12,000 g for 20 minutes at 4°C, and the supernatants were collected in fresh tubes kept on ice.

The protein concentration was determined using the Bradford assay, and subsequently, 40-50 μg of total protein per condition was loaded onto a 12% polyacrylamide gel after boiling the samples in 2X Laemmli sample buffer (4% (w/v) SDS, 20% (v/v) glycerol, 120 mM Tris-HCl pH 6.8, 0.02% bromophenol blue) containing 5% β-mercaptoethanol for 15 minutes. The samples were then immediately resolved on SDS-polyacrylamide gel electrophoresis (PAGE), and the proteins were transferred to Immobilon®-P PVDF membrane (Merck Millipore, #IPVH00010). Immunoblots were blocked with 5% bovine serum albumin (BSA) in tris-buffered saline (TBS) containing 0.1% Tween-20 (TBS-T) for 1 hour at room temperature. The membrane was then probed with specific primary antibodies diluted 1:1000 in antibody dilution buffer (3% BSA in TBS-T) and incubated overnight at 4°C - either anti-TNFR1 rabbit monoclonal antibody (Invitrogen, #MA5-14899) or anti-β-Actin mouse monoclonal antibody (Santa Cruz Biotechnology, #sc-47778). Following primary antibody incubation, the membranes were washed thrice with TBS-T for 5 minutes each and then incubated with the respective secondary antibodies at room temperature for 1 hour - either 1:10,000 horseradish peroxidase (HRP)-conjugated goat anti-rabbit IgG (Sigma-Aldrich/Merck, #A9169) or anti-mouse IgG (Sigma-Aldrich/Merck, #A9044). A standard Pre-stained Protein Ladder (HiMedia, #MBT092) was used to determine the molecular weight of the proteins of interest.

For the detection of GFP, the same procedure was followed, with the exception that the primary antibody used was a monoclonal anti-GFP antibody (Santa Cruz Biotechnology, #sc-9996) at a dilution of 1:1000. For immunoblotting of p65 and phospho-p65, the primary antibodies used were NF-κB p65 (L8F6) Mouse monoclonal antibody (Cell Signaling Technology, #6956) and Phospho- NF-κB p65 (Ser536) (93H1) Rabbit monoclonal antibody (Cell Signaling Technology, #3033), each at a 1:1000 dilution. The corresponding HRP-conjugated secondary antibodies were the same as described before.

Immunodetection was performed using standard chemiluminescence procedures, and densitometric analysis was carried out using ImageJ software.

### Immunoprecipitation

HeLa cells (∼8 × 10⁶) were seeded in 10-cm culture dishes and incubated overnight. Cells were lysed on ice in buffer containing 50 mM Tris-HCl (pH 7.4), 150 mM NaCl, 1% NP-40, 0.1% SDS, and 0.5% sodium deoxycholate, supplemented with protease and phosphatase inhibitor cocktail (Sigma, #PPC1010). Lysates were incubated on ice for 15 min and centrifuged at 15,000 × *g* for 20 min at 4°C to remove debris. Protein concentrations were determined using the Bradford assay.

To minimize nonspecific binding, lysates were precleared with 50 μL of protein A/G–PLUS agarose beads (Santa Cruz, #sc-2003) for 1 hour at 4°C. For immunoprecipitation, 3 mg of precleared lysate was incubated overnight at 4°C with 10 μL of anti-TNFR1/TNFRSF1A rabbit polyclonal antibody (ABclonal, #A1540) and 50 μL of protein A/G–PLUS agarose beads. Immune complexes were recovered by centrifugation and washed three times with lysis buffer before separation by SDS–PAGE on 10% Tris-glycine gels.

Proteins were transferred to PVDF membranes (Millipore) and blocked in TBS containing 0.1% Tween-20 and 5% BSA for 1 hour at room temperature. Membranes were probed overnight at 4°C with rabbit anti–TNF-α antibody (Thermo Fisher, #PA5-19810) or anti–TNFR1/TNFRSF1A antibody (ABclonal, #A1540), each at a 1:1000 dilution. GAPDH served as a loading control and was detected using a mouse monoclonal anti-GAPDH antibody (ABclonal, #AC033). After washing, membranes were incubated with horseradish peroxidase–conjugated secondary antibodies for 1 hour at room temperature: goat anti-rabbit IgG (1:10,000; Sigma-Aldrich, #A0545) and goat anti-mouse IgG (1:10,000; Sigma-Aldrich/Merck, #A9044). Detection was performed using enhanced chemiluminescence.

### Flow cytometry

Before the flow cytometry assay, HeLa cells (1 x 10^5^) were cultured in 35 mm Petri dishes overnight for 16 hours under standard cell culture conditions. Following this, the cells were exposed to various treatments as described in the results section. Following treatment, cells were washed with ice-cold 1× PBS, detached, pelleted in microfuge tubes, and fixed with 4% PFA for 15 minutes on ice. The cells were then washed three times with ice-cold 1× PBS and blocked with 5% BSA in 1× PBS for 20 minutes. Subsequently, they were incubated for 1 hour with TNFR1/TNFRSF1A rabbit pAb (Invitrogen, #PA5-95585; 1:500) prepared in FACS buffer (3% BSA in 1× PBS, 2 mM EDTA). After two washes with ice-cold FACS buffer, cells were incubated with Alexa Fluor 488 conjugated anti-rabbit IgG secondary antibody (Cell Signaling Technology, #4412; 1:200) for 30 minutes, followed by three additional washes with ice-cold FACS buffer. Subsequently, the cells were suspended in FACS buffer to prepare them for flow cytometric analysis. A total of 10^4^ cells were recorded for each condition in the FITC channel to measure the cell surface expression of TNFR1 relative to unstained and control conditions. Data acquisition was performed using a BD FACS Aria III, and the respective fluorescence count histogram and normalized mean fluorescence intensities (MFI) were analyzed using Flowing Software and then plotted using Origin 2019b. The signaling efficacy of TNFR1 was quantified by determining the mean p65 N/C ratio per plasma membrane-associated TNFR1 molecule, calculated using the MFI obtained from flow cytometry in each condition (with or without sTNF-α).

### TNFR1-EGFP chimeric construct validation and cluster area analysis using confocal microscopy

To validate the TNFR1-EGFP chimeric construct, TNFR1 was immunostained in TNFR1-EGFP overexpressing cells without plasma membrane permeabilization as previously stated. The cells were imaged using a Zeiss LSM 780 light scanning confocal system with a 63x oil immersion objective, using two channels for green (TNFR1-EGFP) and red (Alexa Fluor-594, endogenous) fluorescence. Merging the two channels resulted in yellow color in the TNFR1-EGFP cluster regions, indicating colocalization, and confirming that the anti-TNFR1 pAb recognizes our TNFR1-EGFP construct.

For TNFR1 endogenous and exogenous cluster area measurements in different experiments, as stated in the result sections, images were captured using a Zeiss LSM 780 light scanning confocal system with a 63x oil immersion objective in airy scan mode. Then, both endogenous and exogenous (TNFR1-EGFP) clusters were segregated using intensity-based thresholding in ImageJ, and the area of the clusters was measured using the segregated ROI in measurement plugins.

### Quantification of TNFR1-EGFP clusters on the plasma membrane over time

HeLa cells overexpressing TNFR1-EGFP were stained with 0.5x CellMask Orange (CellMask™ Plasma Membrane Stains, Invitrogen, #C10045) at 4-, 12-, 18-, and 24-hours post-transfection. The live HeLa cells overexpressing TNFR1-EGFP and stained with CellMask Orange were imaged using a Zeiss LSM 780 light scanning confocal system. The imaging was performed with a 63x oil immersion objective in airy scan super-resolution mode, with a 0.4μm Z-step size, using two appropriate channels for green (EGFP) and orange (CellMask Orange) fluorescence. Image analysis was performed in ImageJ, where the cell boundary was determined from the CellMask Orange image using intensity-based thresholding. Clusters of TNFR1-EGFP that overlapped with the CellMask Orange-derived cell boundary were classified as plasma membrane-associated clusters. Total and plasma membrane-associated TNFR1-EGFP cluster counting of a cell was conducted using the 3D object counter plugin in ImageJ. The percentage of fractional plasma membrane-associated clusters was then calculated at different time points.

### Nuclear-to-cytoplasmic (N/C) p65 measurement

Fluorescence images of p65-immunostained HeLa cells were captured using a Zeiss AxioObserver Z1 microscope with a 40x air objective to measure the nuclear-to-cytoplasmic ratio (N/C). Z-stacks of randomly selected cell populations were obtained for analysis. Image analysis using ImageJ was performed to determine the N/C ratio of p65. The nuclear portion of p65 was quantified by selecting a region of interest (ROI) in the focused Z-plane of the nucleus, as identified from Hoechst staining images. The mean fluorescence intensity of nuclear p65 (N) was measured using this ROI from the p65 immunostaining z-stack image. For cytoplasmic p65 (C), the total fluorescence intensity of the cytoplasm was measured in areas where the cytoplasm was in focus. The total nuclear fluorescence intensity was then subtracted from this value. The mean cytoplasmic p65 intensity was calculated by dividing the total cytoplasmic intensity by the cytoplasmic area, which was determined by subtracting the nuclear area from the total protoplasmic area.

### Cell culture in a 3D gel-like microenvironment

HeLa cells (1 x 10^5^) were seeded in a glass-bottom Petri dish and allowed to adhere overnight under standard cell culture conditions. Subsequently, the medium was aspirated, and a solution of 1% low-melting HiRes Agarose (Genetix, Puregene, #PG-432) in DMEM was carefully added to cover the adherent cells, forming a thin gel layer. The Petri dish was then briefly incubated at 4°C for 2 minutes to facilitate rapid gel formation. Fresh media was gently added on top of the agarose gel cover to prevent dehydration. The cells were maintained within this 3D gel-like microenvironment for the durations specified in the results section.

### ROS measurements

HeLa cells were seeded in a 96-well plate at equal density (1 x 10^4^ cells) and allowed to grow for 24 hours. Subsequently, the cells were treated with 2 μM cytochalasin D (CytoD) (Sigma-Aldrich/Merck, #C8273) for 1 hour, with or without pre-treatment with 500 μM L-Ascorbic acid (Vitamin C) (Sigma-Aldrich/Merck, #A4544) for 1 hour. For reactive oxygen species (ROS) analysis, cells were stained with 10 µM H_2_DCFDA (Sigma-Aldrich/Merck, #287810) for 30 minutes following each treatment and then lysed with 95% dimethyl sulphoxide (DMSO) (HiMedia, #MB058). The fluorescence intensity of each cell sample was measured using a spectrofluorometer (SpectraMax iD5 Multi-Mode Microplate Reader, Molecular Devices) with excitation/emission wavelengths set at 485 nm/535 nm.

### Fluorescence Anisotropy Imaging Microscopy (FAIM)

Fluorescence anisotropy of TNFR1-EGFP was measured in HeLa cells overexpressing the protein, 18-20 hours post-transfection. The fluorescence anisotropy was determined using a well-established principle (*36*). Imaging was performed with a Zeiss AxioObserver Z1 epifluorescence microscope equipped with a 63x oil immersion objective (NA 1.4) and a mercury arc lamp (HXP 120 V) as the light source. Horizontally polarized excitation light was achieved by passing the light through a linear polarizer (ThorLabs) in the excitation path. In the emission path, a polarizing beam splitter (DV2, Photometrics) was utilized to divide the resulting polarized fluorescence signal into parallel and perpendicular polarizations. The fluorescence from these two polarizations was simultaneously captured by a CMOS camera (Hamamatsu Orca Flash 4.0 C13440), with one half of the image representing the parallel channel (𝐼_‖_) and the other half representing the perpendicular channel (𝐼_⊥_). Due to deformations in the optical path and thermal fluctuations, there was partial overlap between the two halves (𝐼_‖_, 𝐼_⊥_). To correct this, image registration of each cluster of TNFR1-EGFP in the two channels was performed by intensity cross-correlation using the particle image velocimetry (PIV) method, followed by MATLAB’s mono-modal intensity-based registration tool (**fig. S9C**). Subsequently, the fluorescence anisotropy (𝑟) in each pixel of the clusters was calculated using the equation:

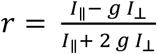

Where *g* is the instrumental correction factor (G-factor, ^𝐼‖^⁄_⊥_), determined by measuring 100 nM fluorescein in water in each pixel. A custom MATLAB code was developed for TNFR1-EGFP cluster image registration and anisotropy calculation. Verification of homo-FRET was carried out through sequential photobleaching at 100% lamp power for 20 seconds, with imaging of the same cell before and after photobleaching. Background correction was performed by subtracting the mean fluorescence intensity of regions without transfected cells in the imaging field. The fluorescence anisotropy of HeLa-mEGFP and HeLa-2GFP cells was assessed using in-house ImageJ code. To achieve image registration, fluorescent polystyrene microspheres with a 200 nm diameter were dried onto a glass coverslip and imaged in the same configuration as the anisotropy measurements. The Descriptor-based Registration plugin of Fiji (ImageJ) was employed for this purpose.

From the anisotropy calculation of TNFR1-EGFP clusters, we obtained a total intensity map (𝐼_‖_ + 2 *g* 𝐼_⊥_) and a fluorescence anisotropy map (𝑟) for each cluster. To simplify our analysis, we divided the fluorescence anisotropy map of each TNFR1-EGFP cluster into two regions, which we named core (central region) and rim (peripheral region), based on the intensity map. In the intensity map, the region with intensity greater than or equal to 70% of the maximum intensity of the cluster was designated as the core, while the region with intensity greater than 10% of the maximum intensity was considered the whole cluster. The rim region was then obtained by subtracting the core region from the whole cluster. Therefore, the rim region comprised the intensity values between >10% and <70% of the maximum intensity of the cluster (**Fig. 1, D and E**). A custom ImageJ code was developed to perform this segmentation. Using the same region of interest (ROI) from the intensity map, we extract the core and rim anisotropy values from the anisotropy map.

### Fluorescence Lifetime Imaging Microscopy (FLIM)

Flipper-TR (Spirochrome, #SC020) was used to assess the plasma membrane tension of cells through fluorescence lifetime imaging. Fluorescence lifetime measurements of Flipper-TR were conducted using an Olympus IX71 microscope with a 60x water immersion objective (NA 1.2) fitted with a time-correlated single-photon counting (TCSPC) module from PicoQuant, and excited with a picosecond 470 nm laser (PicoQuant). The fluorescence lifetime data for each pixel were analyzed by fitting them to a bi-exponential decay curve using SymPhoTime64 software. To quantify the lifetime, the higher amplitude lifetime (𝜏_1_) at each pixel was used, as detailed in the referenced article (*53*). For each cell, 𝜏_1_values were averaged over the region of interest (ROI) corresponding to the plasma membrane. To assess the effect of a 3D gel-like microenvironment on membrane tension, HeLa cells were incubated under a layer of low-melting agarose overnight, while THP-1 cells were incubated for 4 hours. Corresponding 2D control cells were seeded at the same time, 12 hours before agarose overlay. Both 2D (control) and 3D (agarose-covered) cells were simultaneously incubated with 200 nM Flipper-TR for 15–20 minutes before performing all fluorescence lifetime measurements.

### Fluorescence Recovery after Photobleaching (FRAP)

FRAP experiments were conducted on HeLa cells overexpressing TNFR1-EGFP using a Zeiss LSM 780 light scanning confocal system with a 63x oil immersion objective. TNFR1-EGFP clusters were first imaged with 2% laser power and then bleached using 60% laser power at 488 nm, followed by the acquisition of time-lapse imaging for 90 seconds with 1-second intervals using 2% laser power of the same region. The percentage of recovery after bleaching was calculated relative to the pre-bleached condition by comparing the mean fluorescence intensity of the clusters over time to the pre-bleach mean fluorescence intensity. This value was then plotted against time to visualize the recovery process.

### IRM

Cell imaging was conducted on a Nikon Eclipse Ti-E motorized inverted microscope (Nikon, Japan), featuring adjustable field and aperture diaphragms, a 60X Plan Apo lens (NA 1.22, water immersion), and a 1.5X external magnification. The microscope was positioned on an onstage 37°C incubator (Tokai Hit, Japan). Imaging was facilitated using an s-CMOS camera (ORCA Flash 4.0, Hamamatsu, Japan). Illumination was provided by a 100W mercury arc lamp, employing an interference filter (546±12 nm) and a 50-50 beam splitter. For Interference Reflection Microscopy (IRM), movies comprised 2048 frames (20 frames/s), recorded with an exposure time of 50 ms. Analysis of IRM images to obtain fluctuation tension was performed (*72*) using three different steps. At first, the calibration factor is obtained to convert intensity to relative height, and pixels corresponding to the first branch region (FBR) of the interference pattern are identified. Subsequently, the time series of height was obtained for each pixel, and the amplitude of fluctuations (SD_time_ and SD_space_) was obtained for FBRs. Finally, for every FBR, the power spectral density (PSD) was obtained and fitted to an adapted Helfrich-Calham model to extract the fluctuation tension and other parameters (*73*).

### TIRF imaging

TIRF Microscopy utilized an inverted microscope (Olympus IX-83, Olympus, Japan) equipped with a 100X 1.49 NA oil immersion TIRF objective (PlanApo, Olympus). The imaging setup included an s-CMOS camera (ORCA Flash 4.0, Hamamatsu, Japan) and a 488 nm laser source. Image acquisition was performed with an exposure time of 200 ms, achieving a penetration depth of approximately 70 nm. For cluster size analysis from TIRF images in different conditions, as depicted in the result sections, local thresholding was performed to detect objects using MATLAB. All immobile clusters were manually identified, and the area of the objects from the region properties was used for comparison.

Time-lapse imaging was performed in TIRF mode on TNFR1-EGFP–overexpressing HeLa cells (**fig. S1C**) using continuous acquisition with a 50-ms exposure for 5 minutes, without intervals, to assess receptor cluster stoichiometry.

### Statistical analysis

Individual measurements from technical replicates within each treatment group were pooled to generate a unified dataset of biological replicates. Variations in mean values of measurements taken from the same cells before and after treatment were assessed using a paired two-tailed Student’s *t*-test. For comparisons between two distinct cell populations, an unpaired two-tailed Student’s *t*-test was applied, assuming that sufficiently large sample sizes approximate a normal distribution. When comparisons involved more than two groups, a one-way ANOVA was performed. Details of the statistical tests applied are provided in the respective figure legends. All statistical analyses and graphical representations were performed using Origin 2019b.

## Acknowledgments

We express our gratitude to the Central Scientific Services (CSS) of the Indian Association for the Cultivation of Science (IACS), Kolkata, for providing us with access to instrumental facilities. Special thanks to Mrs. Debapriya Ghatak, Mr. Chanchal Kumar Das, and Mr. Gopal Krishna Manna for their invaluable assistance with confocal microscopy, FACS, and graphical illustration, respectively. We are indebted to Prof. Maria Vartiainen (University of Helsinki, Finland) for generously gifting us with the GFP-GFP (2GFP) plasmid. Our appreciation goes to Dr. Dipyaman Ganguly (Indian Institute of Chemical Biology, Kolkata) for graciously allowing us to utilize FACS and to his student, Ms. Shreya Roy, for assisting with flow cytometry data acquisition. We also extend our gratitude to Dr. Prosenjit Sen (IACS, Kolkata) for providing us with an essential cell line (THP-1). BS thanks Wellcome Trust/DBT India Alliance fellowship (Grant Number IA/I/13/1/500885) for the TIRF imaging. We acknowledge ChatGPT 3.5 for rephrasing our write-up for clarity. Finally, we thank Dr. Mahesh Agarwal for critically reading the manuscript and providing valuable suggestions, which greatly contributed to preparing this manuscript for publication. Furthermore, we acknowledge IACS, Kolkata, for providing intramural funds.

## Funding

SERB Core Research Grant, Department of Science and Technology, Ministry of Science and Technology CRG/2022/005356 (DKS)

Department of Biotechnology, Ministry of Science and Technology BT/PR6995/BRB/10/1140/2012 (DKS)

SERB Grant number SERB_CRG_2458 (BS) CEFIPRA Grant number 6303-1 (BS)

Indian Association for the Cultivation of Science, Kolkata 700032, India (SJ, PR, PB, BBB, NN)

Council of Scientific and Industrial Research 09/080(1074)/2018-EMR-I (PR) University Grants Commission 201610161681 (JD)

## Author contributions

SJ and PR are co-first authors of this paper and contributed equally. DKS, BS, SJ, and PR designed the project and prepared the manuscript. SJ, PR, and JD conducted the experiments and analyzed the data. JD performed the IRM and TIRF imaging. PB was responsible for setting up and calibrating the HeLa-mEGFP and HeLa-2GFP fluorescence anisotropy measurements. BBB and NN contributed to data analysis. BS assisted in reviewing the manuscript.

## Data and materials availability

All data are available in the main text or the supplementary materials. Codes used in the manuscript are freely available online (<u>https:/github.com/SubhamoyJana/TNFR1-Project</u>).

