## Supplementary Materials for "Intra-cluster receptor density (IRD): a molecular switch for TNFR1 clusters’ signaling"

Subhamoy Jana *et al.*

*Corresponding author: Deepak Kumar Sinha


**This PDF file includes:**

Figs. S1 to S9
Table S1
Movies S1 to S3

**Other Supplementary Materials for this manuscript include the following:**

Movies S1 to S3

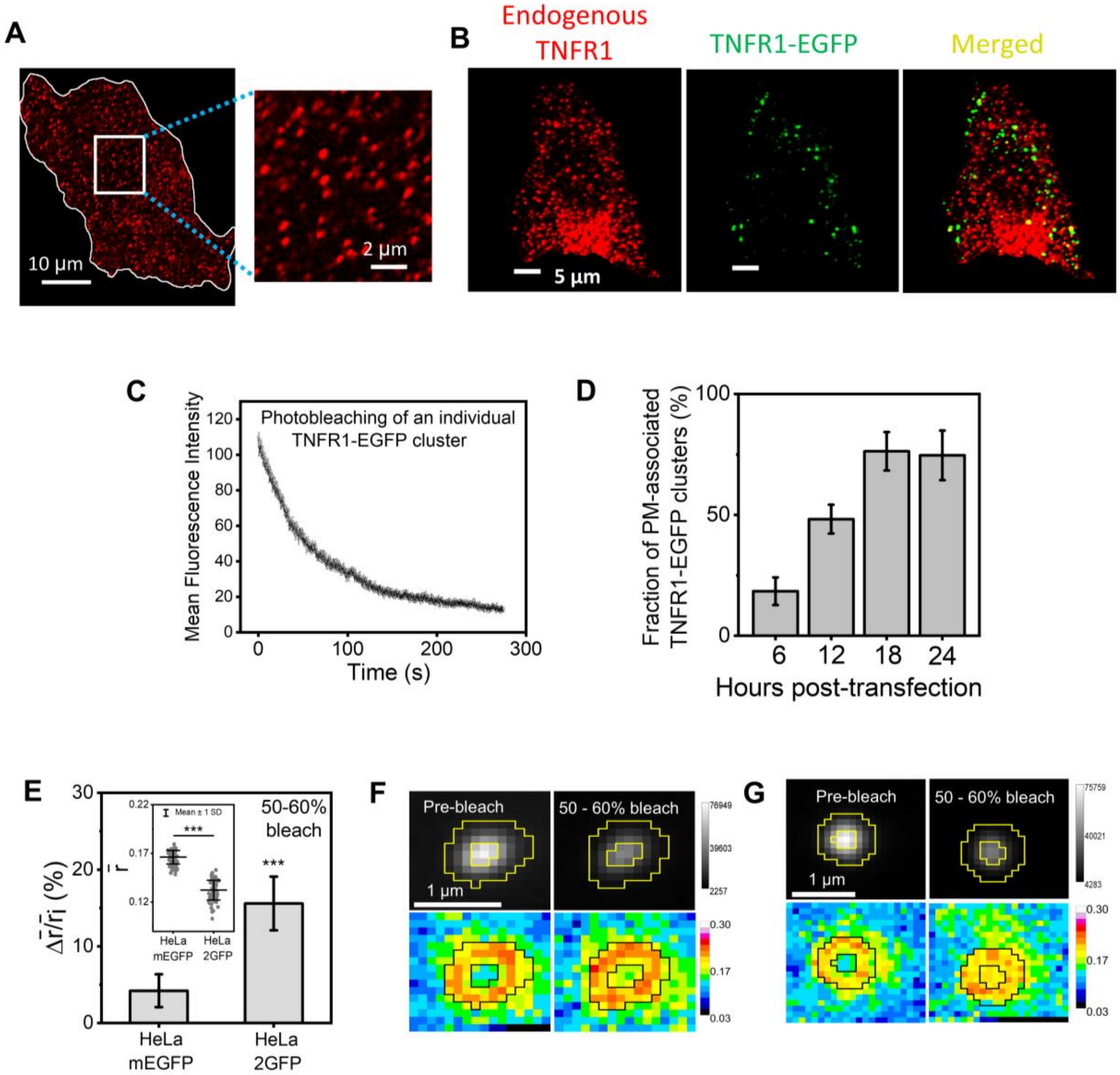


**Fig. S1. Homo-FRET analysis of TNFR1-EGFP clusters reveals spatial organization and dynamic remodeling of TNFR1 in HeLa cells.**

(**A**) Representative confocal Airyscan image of a HeLa cell immunostained with anti-TNFR1 antibody (red, left), with a magnified view highlighting endogenous TNFR1 clusters (right). (**B**) Representative confocal images of a HeLa cell expressing TNFR1-EGFP (green, middle) and immunostained with anti-TNFR1 antibody (red, left); merged image (right) shows their colocalization. (**C**) Mean fluorescence intensity of a single TNFR1-EGFP cluster plotted over time during photobleaching. (**D**) Quantification of the fraction of plasma membrane (PM)–associated TNFR1-EGFP clusters (number of PM clusters/total number of clusters) over time posttransfection. Data represent means ± SD from 3 cells each in two independent experiments. (**E**) Percentage change in relative mean fluorescence anisotropy ($\Delta\bar{r}/\bar{r}_i$) of mEGFP and 2EGFP overexpressed HeLa cells (n = 70) upon 50-60% photobleaching. Data are presented as means ± SD from 10 cells each in three independent experiments. Here, $\Delta\bar{r} = \bar{r}_f - \bar{r}_i$ where $\bar{r}_f$ and $\bar{r}_i$ represent post- and pre-bleach mean anisotropy, respectively. The inset shows the mean fluorescence anisotropy ($\bar{r}$) for each construct across three independent experiments. Since 2EGFP forms a close dimer of mEGFP, homo-FRET occurs within the dimer, leading to an increase in fluorescence anisotropy upon photobleaching. (**F**) Representative fluorescence intensity (top) and corresponding fluorescence anisotropy (bottom) maps of a TNFR1-EGFP cluster before and after photobleaching. (**G**) Representative TNFR1-EGFP cluster showing fluorescence intensity (top) and corresponding fluorescence anisotropy (bottom) maps before and after photobleaching, illustrating uniform fluorescence anisotropy across both the core and the rim regions of the cluster upon photobleaching. Statistical significance was determined by an unpaired two-tailed Student's *t*-test (E) and is indicated in the figures as follows: ∗P < 0.05; ∗∗P < 0.01; ∗∗∗P < 0.001; ns, not significant. Scale bars are indicated in the corresponding images.

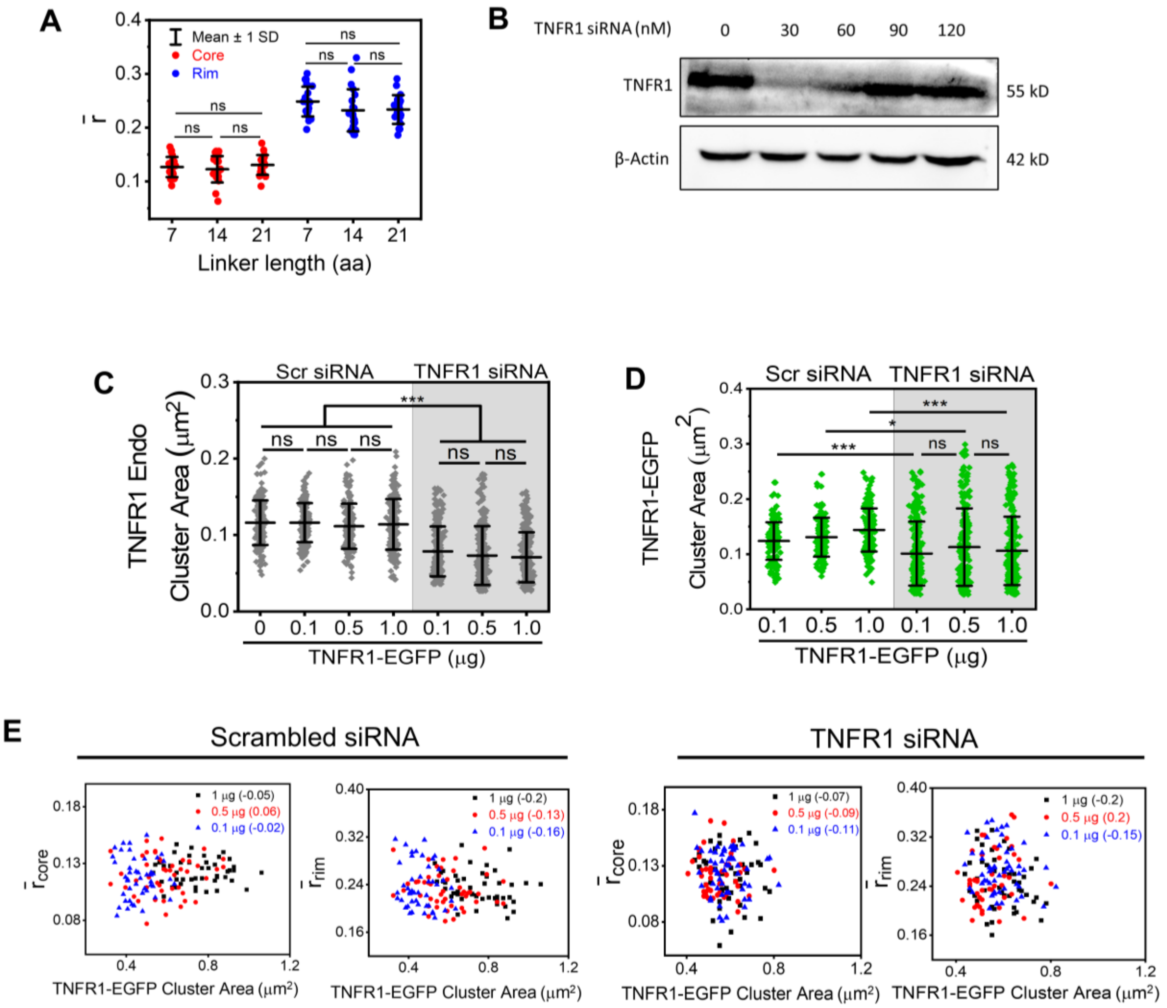


**Fig. S2. TNFR1 expression level and IRD influence cluster area and organization.**

(**A**) Mean fluorescence anisotropy ($\bar{r}$) of the core (red) and rim (blue) regions of TNFR1-EGFP clusters (n = 21) in HeLa cells expressing TNFR1-EGFP constructs with different linker lengths (7, 14, or 21 amino acids). The data represent 2 cells each from two independent experiments. (**B**) Immunoblot showing TNFR1 expression in HeLa cells treated with increasing doses of TNFR1 siRNA. β-Actin was used as a loading control. Representative of two independent experiments. (**C**) Quantification of the area (μm²) of endogenous TNFR1 clusters (n ≥ 170) in TNFR1-EGFP–transfected HeLa cells pre-treated with scrambled siRNA (Scr siRNA) or TNFR1 siRNA, as acquired by confocal Airyscan imaging. The data represent 8 cells each from three independent experiments. (**D**) Quantification of the area (μm²) of TNFR1-EGFP clusters (n ≥ 170) in TNFR1-

EGFP–transfected HeLa cells pre-treated with scrambled siRNA (Scr siRNA) or TNFR1 siRNA, as acquired by confocal Airyscan imaging. The data represent 8 cells each from three independent experiments. (**E**) Scatter plots showing the relationship between TNFR1-EGFP cluster area (μm$^2$) and mean fluorescence anisotropy of the core ($\bar{\boldsymbol{r}}_{\boldsymbol{core}}$) and rim ($\bar{\boldsymbol{r}}_{\boldsymbol{rim}}$) of TNFR1-EGFP clusters (n = 45) in HeLa cells transfected with scrambled siRNA (left) or TNFR1 siRNA (right), and co-transfected with increasing doses (0-1 μg) of TNFR1-EGFP plasmid. Data are representative of 3 cells each in three independent experiments. Pearson's correlation coefficients are indicated in the figure for the corresponding TNFR1-EGFP doses. Statistical significance was determined by one-way ANOVA (A, C, D) and is indicated in the figures as follows: $*P < 0.05$; $**P < 0.01$; $***P < 0.001$; ns, not significant. Scale bars are indicated in the corresponding images.

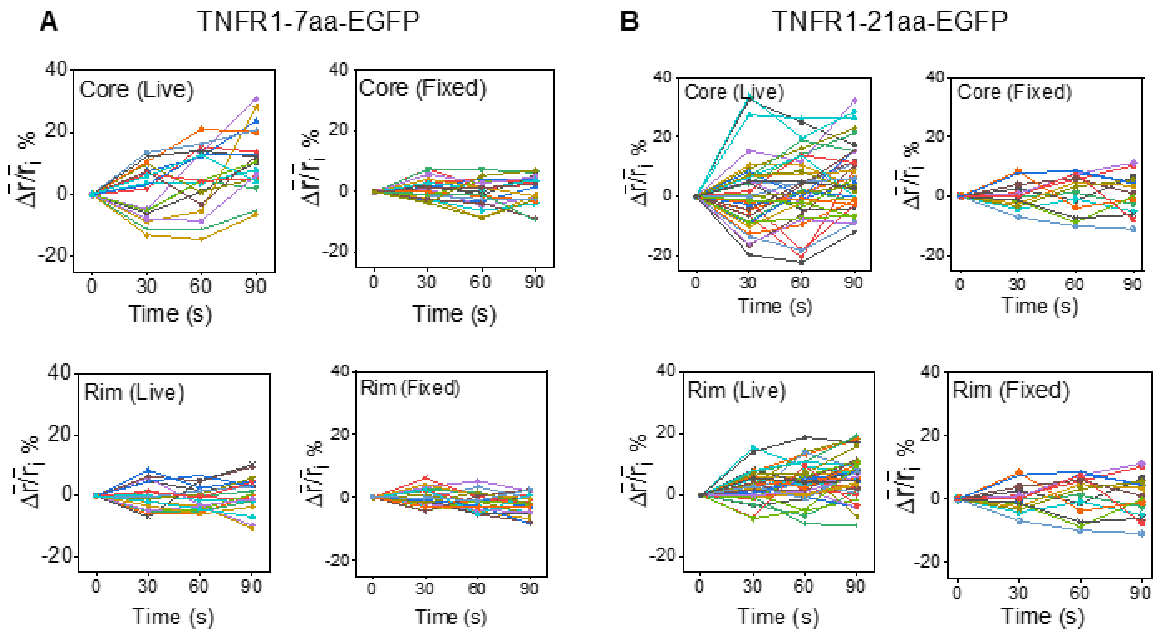


**Fig. S3. Temporal dynamics of fluorescence anisotropy in live and fixed TNFR1-EGFP clusters with variable linker lengths.**

**(A)** Percentage change in relative mean fluorescence anisotropy ($\Delta\bar{\boldsymbol{r}}/\bar{\boldsymbol{r}}_{\boldsymbol{i}}$), where $\Delta\bar{\boldsymbol{r}} = \bar{\boldsymbol{r}}(\boldsymbol{t}) - \bar{\boldsymbol{r}}_{\boldsymbol{i}}$, for the core (top) and rim (bottom) regions plotted as a function of time ($\boldsymbol{t}$) in live (left, n = 18) and fixed (right, n = 22) HeLa cells expressing TNFR1-7aa-EGFP, acquired over a 90-second time course. Here, $\bar{\boldsymbol{r}}_{\boldsymbol{i}}$ represents the initial mean anisotropy ($\boldsymbol{t}$ = 0), and $\bar{\boldsymbol{r}}(\boldsymbol{t})$ denotes the mean anisotropy at time $\boldsymbol{t}$. The data represent 2 cells each from two independent experiments. **(B)** Percentage change in relative mean fluorescence anisotropy ($\Delta\bar{\boldsymbol{r}}/\bar{\boldsymbol{r}}_{\boldsymbol{i}}$) for the core (top) and rim (bottom) regions plotted as a function of time ($\boldsymbol{t}$) in live (left, n = 35) and fixed (right, n = 15) HeLa cells expressing TNFR1-21aa-EGFP, acquired over a 90-second time course. The data represent 2 cells each from two independent experiments.

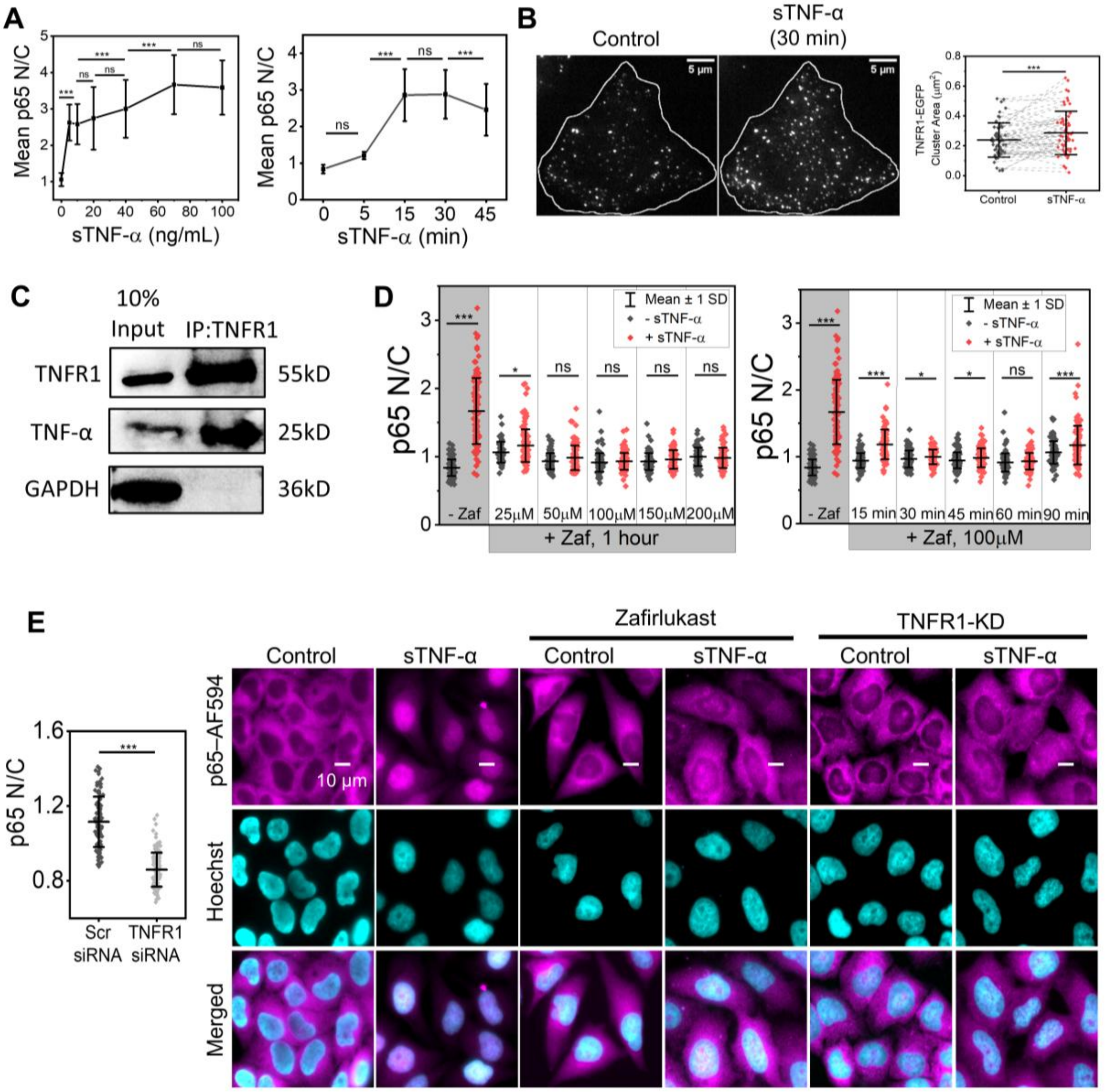


**Fig. S4. Differential modulation of TNFR1-mediated NF-κB activation by sTNF-α, zafirlukast, and receptor expression levels.**

(**A**) Quantification of the nuclear-to-cytoplasmic (N/C) mean fluorescence intensity ratio of p65 in HeLa cells treated with increasing concentrations of sTNF-α for 30 min (left), and with 20 ng/mL sTNF-α for varying durations (right). Data represent mean ± SD from 30 cells per condition across three independent experiments. (**B**) Representative TIRF images of TNFR1-EGFP-expressing HeLa cells before (Control) and after stimulation with 20 ng/mL sTNF-α for 30 min. Quantification of the area (µm²) of the same TNFR1-EGFP clusters (n = 60) before (Control) and after stimulation is shown. The data represent 2 cells each from three independent experiments. (**C**) Immunoblot analysis of TNFR1 and TNF-α in HeLa cell lysates subjected to immunoprecipitation (IP) using an anti-TNFR1 antibody. Both input and IP:TNFR1 samples show positive bands for TNFR1 and TNF-α, while GAPDH served as a negative control. Representative of two independent experiments. (**D**) Quantification of p65 N/C ratio in HeLa cells treated with increasing concentrations of zafirlukast (Zaf) for 1 h in the presence or absence of sTNF-α (left), and in cells treated with 100 μM Zaf for varying durations with or without sTNF-α stimulation (right). The data represent 30 cells per condition across three independent experiments. (**E**) Quantification of p65 N/C mean fluorescence intensity ratio in HeLa cells transfected with scrambled (Scr) or TNFR1 siRNA (left). Data represent mean ± SD from 30 cells per condition across three independent experiments. Representative immunofluorescence images (right) show p65 (magenta, top) in HeLa cells treated with sTNF-α (20 ng/mL, 30 min) under control, zafirlukast-treated (100 μM, 1 h), and TNFR1-knockdown (TNFR1-KD) conditions. DNA was counterstained with Hoechst 33342 (cyan, middle); merged images are shown in the lower panel. Statistical significance was determined using one-way ANOVA (A), paired two-tailed Student's *t*-test (B), or unpaired two-tailed Student's *t*-test (C, D), and is indicated in the figures as follows: $*P < 0.05$; $**P < 0.01$; $***P < 0.001$; ns, not significant. Scale bars are indicated in the corresponding images.

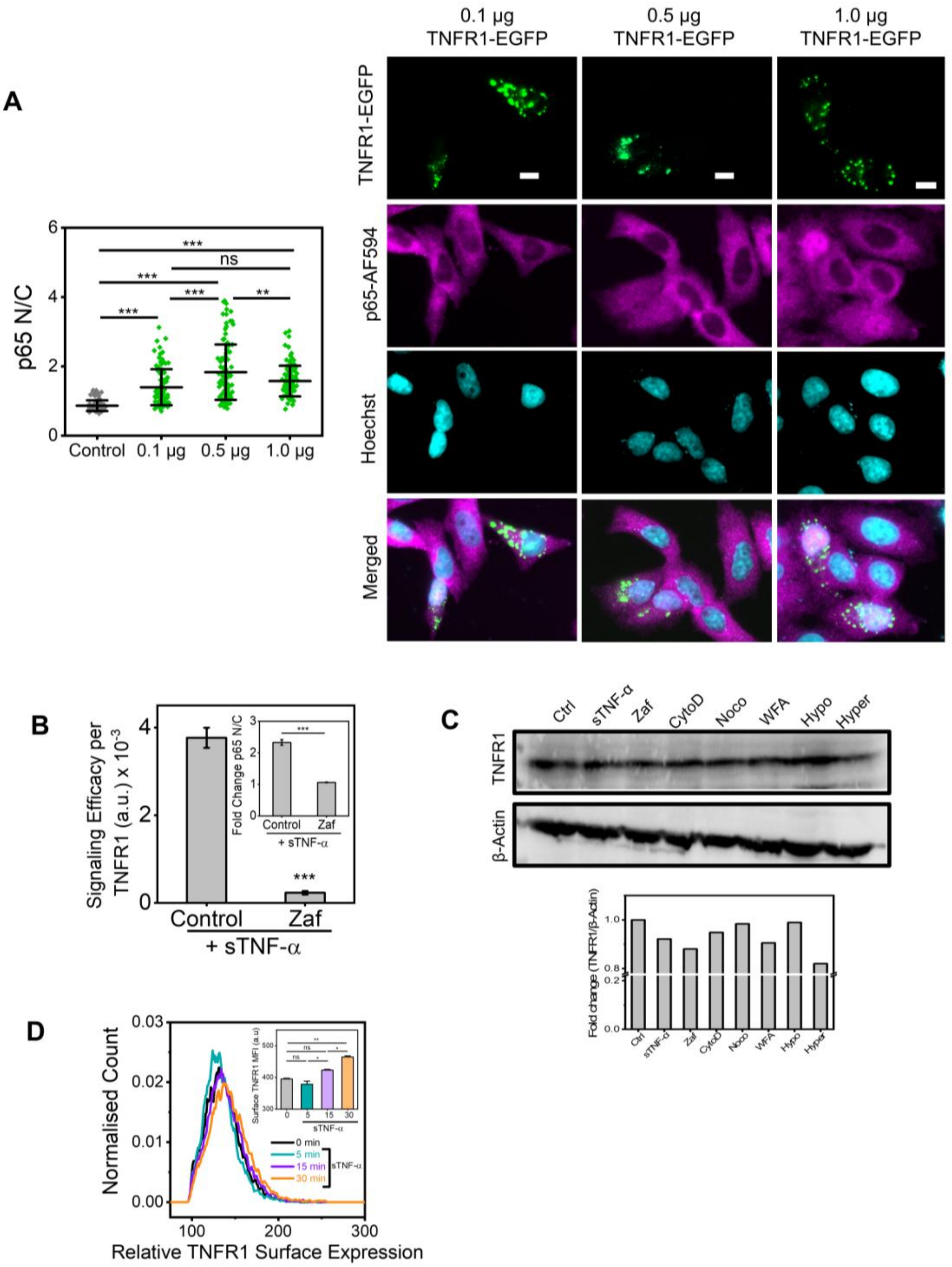


**Fig. S5. Pharmacological perturbations and TNFR1 expression levels modulate signaling efficacy and receptor dynamics.**

(**A**) Quantification of the nuclear-to-cytoplasmic (N/C) mean fluorescence intensity ratio of p65 in HeLa cells transfected with increasing doses (0-1 µg) of TNFR1-EGFP (left). Data represent mean ± SD from 30 cells per condition across three independent experiments. Representative immunofluorescence images (right) show TNFR1-EGFP (green, top), p65 (magenta, second), nuclei stained with Hoechst 33342 (cyan, third), and merged channels (bottom). (**B**) Signaling efficacy of TNFR1 in control and zafirlukast-treated (Zaf) HeLa cells in response to sTNF-α. Signaling efficacy represents the change in N/C ratio of p65 normalized to surface TNFR1 levels (ΔN/C p65 ratio / respective surface TNFR1 MFI). The inset shows fold changes in p65 N/C ratio normalized to their respective basal (unstimulated) levels. Data represent mean ± SD from 30 cells per condition across three independent experiments. (**C**) Immunoblot analysis of total TNFR1 protein levels in HeLa cells under various treatments — control (Ctrl), sTNF-α (20 ng/mL, 5 min), zafirlukast (Zaf, 100 μM, 1 h), cytochalasin D (CytoD, 2 μM, 1 h), nocodazole (Noco, 18 μM, 2 h), withaferin A (WFA, 2 μM, 4 h), hypotonic (Hypo, 75% $H_2O$, 2 min), and hypertonic (Hyper, 150 mM mannitol, 10 min) conditions. β-Actin served as a loading control. Densitometric quantification of fold change in TNFR1 expression normalized to β-Actin is shown below. (**D**) Flow cytometric analysis of surface TNFR1 expression in control (0 min) and sTNF-α-treated (20 ng/mL, 5–30 min) HeLa cells. Quantification of median fluorescence intensity (MFI) for each condition is shown in the inset. The data represent two independent experiments. Statistical significance was determined using one-way ANOVA (A, D) or unpaired two-tailed Student's *t*-test (B) and is indicated in the figures as follows: ∗$P < 0.05$; ∗∗$P < 0.01$; ∗∗∗$P < 0.001$; ns, not significant. Scale bars are indicated in the corresponding images.

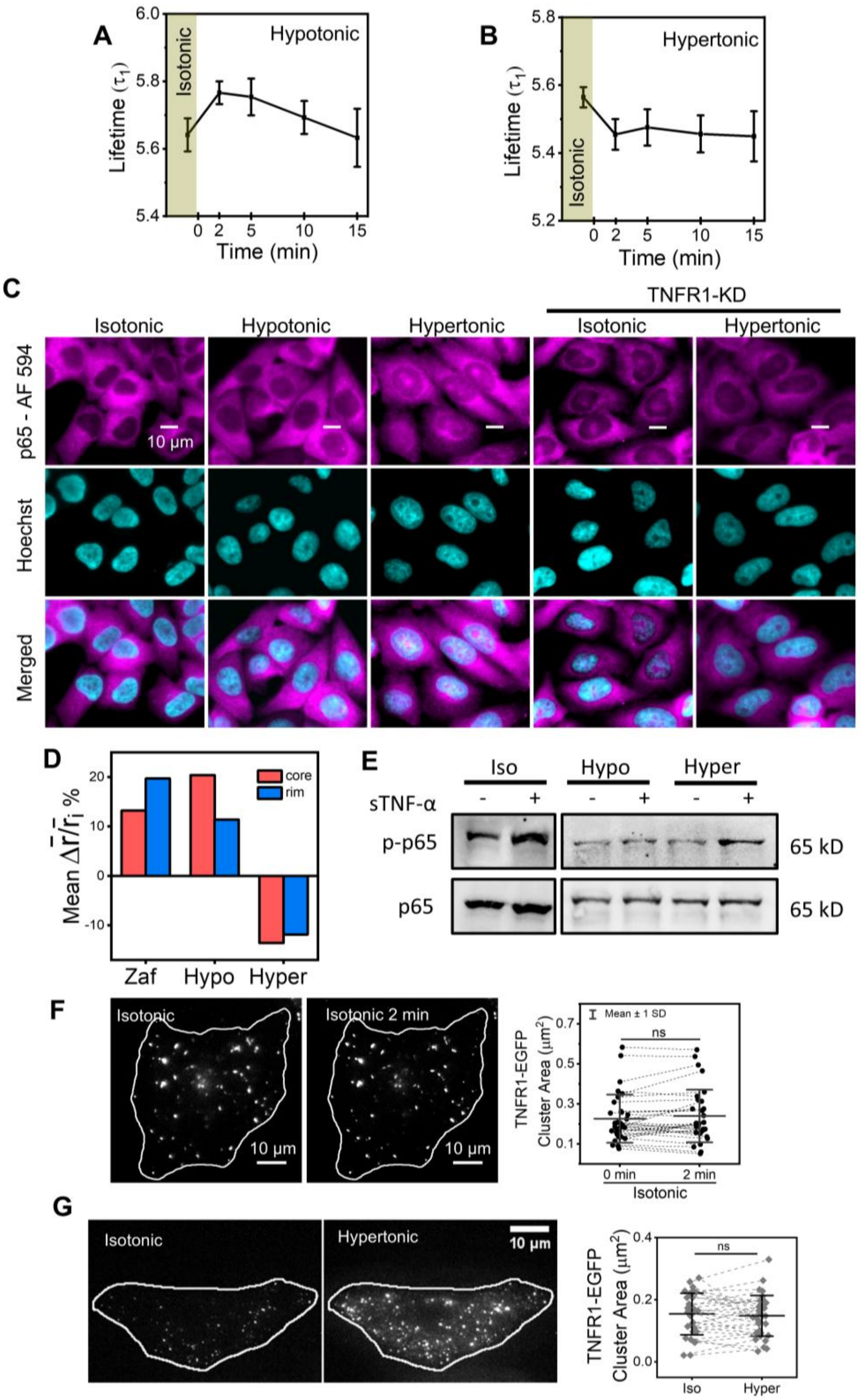


**Fig. S6. Membrane tension regulates TNFR1 IRD and downstream NF-κB signaling in HeLa cells.**

(**A** and **B**) Fluorescence lifetime ($\tau_1$) measurements of the membrane tension probe Flipper-TR in HeLa cells under hypotonic (A) and hypertonic (B) conditions over 15 minutes. Data represent mean ± SD from 5 cells each across two independent experiments. (**C**) Representative immunofluorescence images of HeLa cells showing NF-κB p65 localization (magenta, top) under isotonic, hypotonic, and hypertonic conditions in control and TNFR1-knockdown (TNFR1-KD) cells. Nuclei were counterstained with Hoechst 33342 (cyan, middle), and merged images are shown in the bottom panel. (**D**) Percentage change in relative mean fluorescence anisotropy ($\Delta\,\bar{\boldsymbol{r}}/\bar{\boldsymbol{r}}_{\boldsymbol{i}}$) of the core (red) and rim (blue) regions of TNFR1-EGFP clusters (n = 60) in HeLa cells treated with zafirlukast (Zaf), hypotonic (Hypo), or hypertonic (Hyper) conditions, normalized to respective controls. Here, $\Delta\bar{\boldsymbol{r}} = \bar{\boldsymbol{r}}_{\boldsymbol{f}} - \bar{\boldsymbol{r}}_{\boldsymbol{i}}$ where $\bar{\boldsymbol{r}}_{\boldsymbol{f}}$ and $\bar{\boldsymbol{r}}_{\boldsymbol{i}}$ represent the final and initial mean anisotropy for each treatment, respectively. The data represent 5 cells each across three independent experiments. (**E**) Immunoblot analysis of Ser536 phosphorylated p65 (p-p65) levels in HeLa cells subjected to hypotonic (Hypo) or hypertonic (Hyper) conditions, followed by sTNF-α stimulation, compared to isotonic controls (Iso). Total p65 served as a loading control. Representative of two independent experiments. (**F**) Representative TIRF images of TNFR1-EGFP–expressing HeLa cells under isotonic conditions acquired at 2-minute intervals (left). Quantification of the area ($\mu m^2$) of the same TNFR1-EGFP clusters (n = 32) before and after 2 minutes is shown (right). The data represent 2 cells each in two independent experiments. (**G**) Representative TIRF images of TNFR1-EGFP–transfected HeLa cells before and after 10 minutes of hypertonic shock (left). Quantification of the area ($\mu m^2$) of the same TNFR1-EGFP clusters (n = 34) before (Iso) and after (Hyper) hyperosmotic shock is shown (right). The data represent 2 cells each in two independent experiments. Statistical significance was determined by paired two-tailed Student's *t*-test (F, G) and is indicated in the figures as follows: ∗P < 0.05; ∗∗P < 0.01; ∗∗∗P < 0.001; ns, not significant. Scale bars are indicated in the corresponding images.

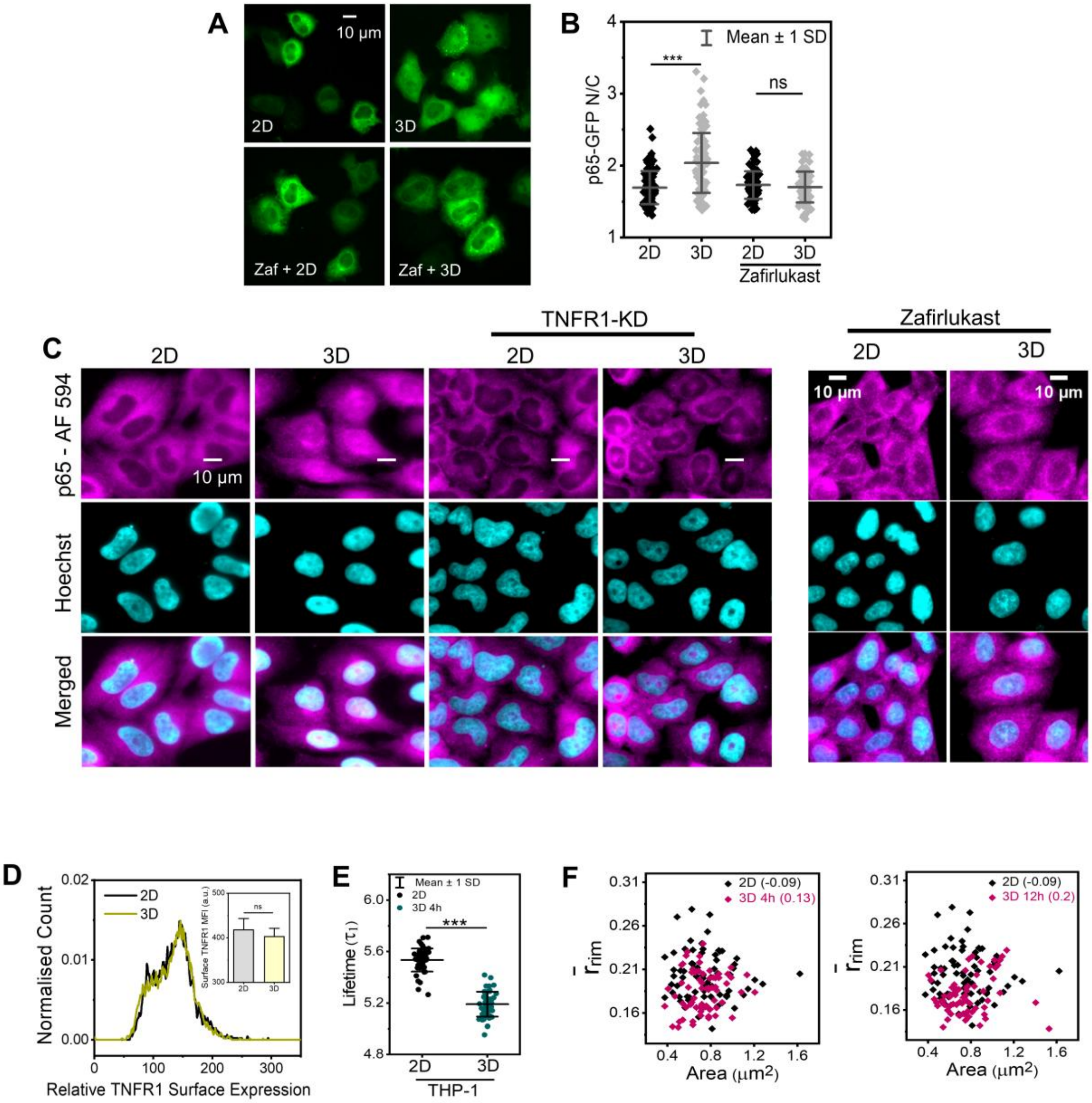


**Fig. S7. TNFR1 signaling and membrane tension are modulated by a 3D gel-like microenvironment.**

(**A**) Live-cell imaging of representative HeLa cells expressing NF-κB–GFP (p65-GFP) cultured in 2D (control) and 3D (1% agarose gel overlay, 12 h) under untreated and zafirlukast-treated (Zaf) conditions. (**B**) Quantification of nuclear-to-cytoplasmic (N/C) mean fluorescence intensity ratio of p65-GFP in live HeLa cells under the indicated conditions. The data represent 35 cells per condition across three independent experiments. (**C**) Representative immunofluorescence images of p65 (magenta, top) in HeLa cells cultured in 2D or 3D under control, TNFR1-knockdown (TNFR1-KD), and zafirlukast-treated conditions. Nuclei were counterstained with Hoechst 33342 (cyan, middle), and merged images are shown in the bottom panel. (**D**) Surface expression of endogenous TNFR1 in HeLa cells cultured in 2D and 3D conditions, analyzed by flow cytometry. The inset shows the median fluorescence intensity (MFI) for the 2D and 3D conditions. (**E**) Fluorescence lifetime ($\tau_1$) of the membrane tension probe Flipper-TR in THP-1 cells cultured in 2D and 3D conditions. The data represent 16 cells per condition across three independent experiments. (**F**) Scatter plots showing TNFR1-EGFP cluster area ($\mu m^2$) versus mean fluorescence anisotropy of the rim ($\bar{\boldsymbol{r}}_{\boldsymbol{rim}}$) of TNFR1-EGFP clusters (n = 70) in HeLa cells cultured in 2D and 3D conditions [4 h (left) and 12 h (right)]. Pearson's correlation coefficients are indicated for each condition. The data represent 3 cells per condition in three independent experiments. Statistical significance was determined by one-way ANOVA (B) or unpaired two-tailed Student's *t*-test (D, E), as indicated in the figures as follows: $*P < 0.05$; $**P < 0.01$; $***P < 0.001$; ns, not significant. Scale bars are indicated in the corresponding images.

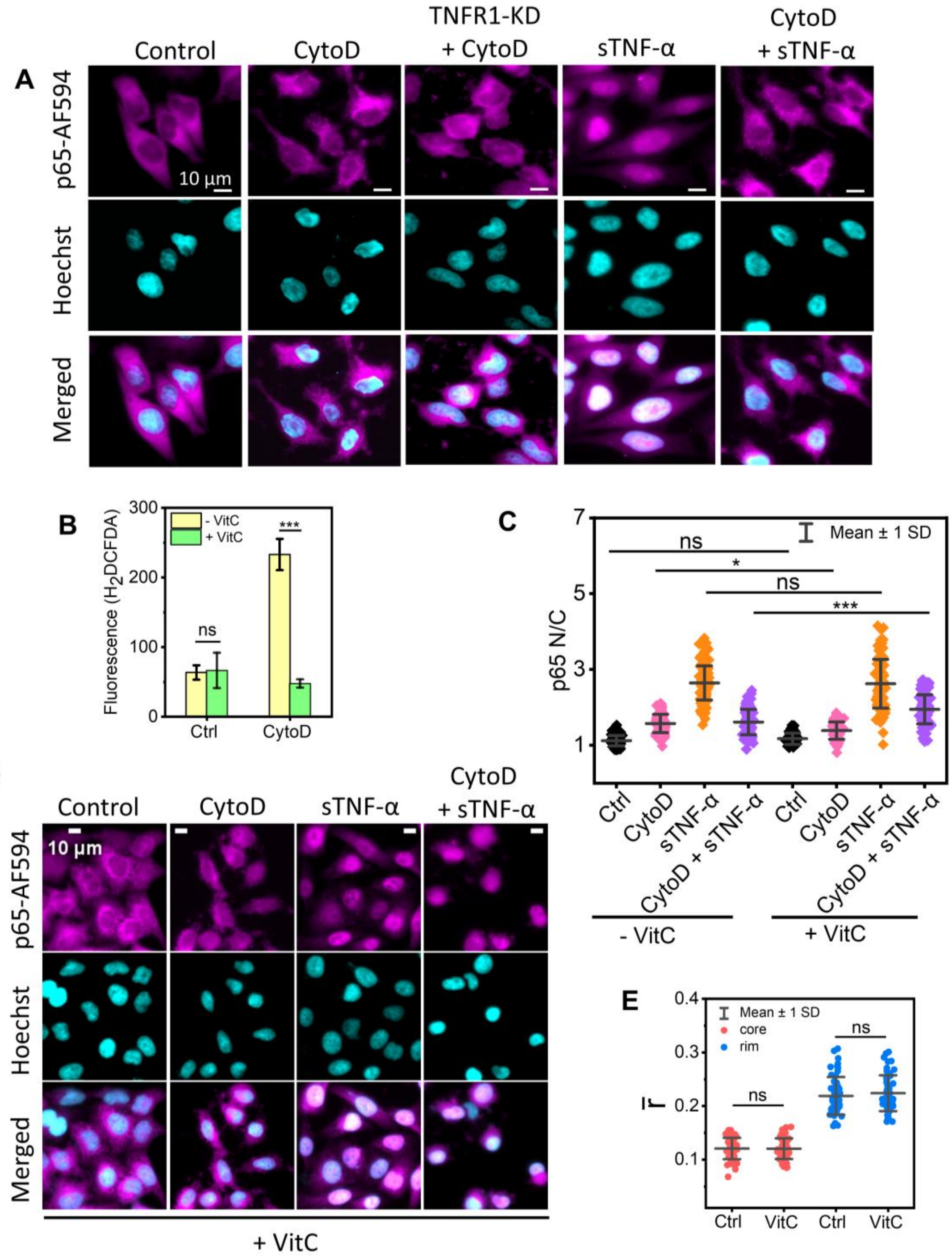


**Fig. S8. Impact of actin disruption on TNFR1 signaling and cluster organization.**

(**A**) Representative immunofluorescence images of HeLa cells stained for p65 (magenta, top) under cytochalasin D (CytoD) treatment in control and TNFR1-knockdown (TNFR1-KD) cells, and following sTNF-α stimulation in the presence or absence of CytoD. Nuclei were counterstained with Hoechst 33342 (cyan, middle), and merged images are shown in the bottom panel. (**B**) Quantification of mean $H_2$DCFDA fluorescence intensity (a.u.) in control (Ctrl) and CytoD-treated HeLa cells, with or without Vitamin C (VitC) pre-treatment. (**C**) Nuclear-to-cytoplasmic (N/C) mean fluorescence intensity ratio of NF-κB (p65) in HeLa cells analyzed by immunostaining under control (Ctrl), CytoD-treated, sTNF-α-treated, and combined CytoD + sTNF-α-treated conditions, with or without VitC pre-treatment. The data represent mean ± SD from 35 cells per condition from three independent experiments. (**D**) Representative immunofluorescence images of p65 (magenta, top) in HeLa cells pre-treated with VitC under control, CytoD-treated, sTNF-α-treated, and combined CytoD + sTNF-α-treated conditions. Nuclei were counterstained with Hoechst 33342 (cyan, middle), and merged images are shown in the bottom panel. (**E**) Mean fluorescence anisotropy ($\bar{\boldsymbol{r}}$) of the core (red) and rim (blue) regions of TNFR1-EGFP clusters (n = 53) in control (Ctrl) and VitC-treated HeLa cells. Data represent 3 cells per condition across three independent experiments. Statistical significance between datasets was determined by unpaired, two-tailed Student's *t*-test (B, E) or one-way ANOVA (C) and is indicated in the figures as follows: $*P < 0.05$; $**P < 0.01$; $***P < 0.001$; ns, not significant. Scale bars are indicated in the corresponding images.

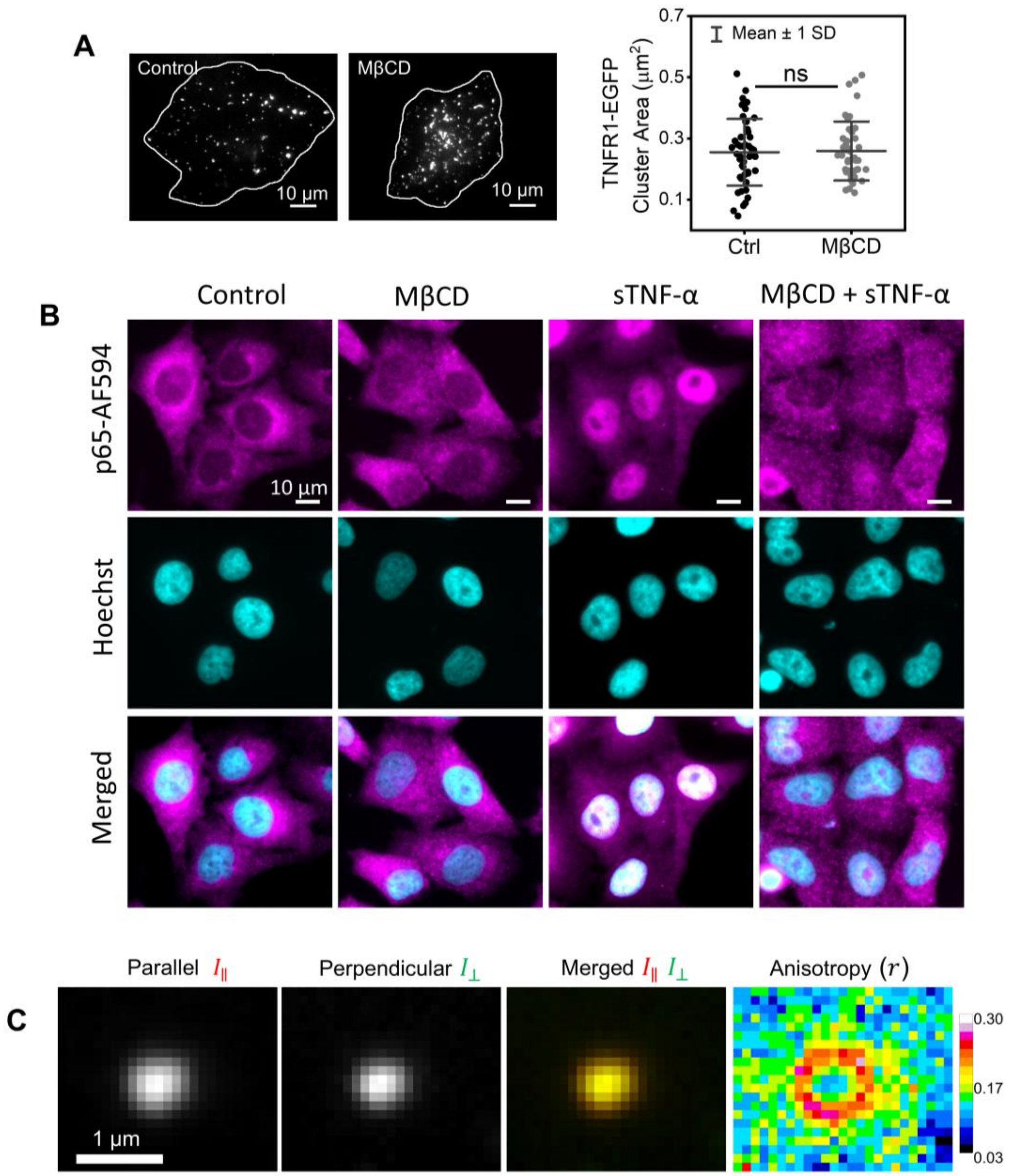


**Fig. S9. Effects of membrane cholesterol depletion on TNFR1 clustering and signaling.**

(**A**) Representative TIRF images (left) of a TNFR1–EGFP–expressing HeLa cell before and after treatment with methyl-β-cyclodextrin (MβCD, 2 mM, 1 hour). Quantification of immobile TNFR1–EGFP cluster area (µm²) before (Ctrl, n = 49) and after (n = 43) MβCD treatment is shown (right). Data represent means ± SD from 3 cells across two independent experiments. (**B**) Representative immunofluorescence images of p65 (magenta, top) in HeLa cells under control, MβCD-treated, sTNF-α-treated, and combined MβCD + sTNF-α-treated conditions. Nuclei were counterstained with Hoechst 33342 (cyan, middle), and merged images are shown in the bottom panel. (**C**) Representative epifluorescence images of a TNFR1–EGFP cluster in parallel (left) and perpendicular (second left) polarization channels, with the merged image (second right) and the corresponding anisotropy map (right). Statistical significance was determined by unpaired two-tailed Student's *t*-test (A) and is indicated in the figures as follows: ∗$P < 0.05$; ∗∗$P < 0.01$; ∗∗∗$P < 0.001$; ns, not significant. Scale bars are indicated in the corresponding images.

**Table S1. Summary of inhibitors that modulate TNFR1 clustering or signaling, along with their mechanisms of action and corresponding references.**

| Inhibitor | Target / Mode of Action | Reference |
|---|---|---|
| **SPD304** | Binds to TNF-α and dissociates its trimeric structure, thus preventing TNFR1 activation. | (74, 75) |
| **IW927** | Directly binds TNFR1 and blocks TNF-α binding, thereby disrupting receptor activation. | (76) |
| **Zafirlukast** | Inhibits PLAD-mediated dimerization of TNFR1, thus impairing receptor clustering and signaling. | (33, 34) |
| **UCB-9260** | Stabilizes an asymmetric TNF-α trimer, preventing effective TNFR1 engagement for downstream signal activation. | (77) |
| **F002** | Cavity-induced allosteric modifier (CIAM) that prevents TNF-α binding to TNFR1 and thereby blocks subsequent pathway activation. | (*78*) |
| **SGT11, C7** | Bind to TNFR1 allosteric sites; inhibit TNF-α–induced NF-κB signaling. | (79) |
| **Suramin, Evans Blue, Trypan Blue** | Interfere with TNF-α trimerization, indirectly preventing TNFR1 activation. | (80, 81) |
| **Cyclic peptides** | Selectively bind TNFR1 and inhibit receptor-ligand interaction. | (82) |

**Movie S1.**

3D-rendered images of a HeLa cell viewed from different orientations, showing the distribution of TNFR1-EGFP clusters (green). The plasma membrane is stained with CellMask Orange (orange).

**Movie S2.**

Time-lapse imaging of TNFR1-EGFP clusters in a HeLa cell, comprising 20 frames captured at 2-second intervals. The movie shows intracellular mobile clusters (yellow circle) and plasma membrane–associated clusters that remain relatively immobile (red circle).

**Movie S3.**

Time-lapse imaging of TNFR1-EGFP clusters in untreated control (left) and dynasore-treated (80 μM, 50 min, right) HeLa cells. Imaging was performed continuously for 600 frames with an acquisition rate of 200 ms per frame.
